# Rapid nucleus-scale reorganization of chromatin in neurons enables transcriptional adaptation for memory consolidation

**DOI:** 10.1101/2020.12.03.409623

**Authors:** Manuel Peter, Dominik F. Aschauer, Renata Vaz Pandolfo, Anne Sinning, Florian Grössl, Dominic Kargl, Klaus Kraitsy, Thomas R. Burkard, Heiko J. Luhmann, Wulf Haubensak, Simon Rumpel

## Abstract

The interphase nucleus is functionally organized in active and repressed territories defining the transcriptional status of the cell. However, it remains poorly understood how the nuclear architecture of neurons adapts in response to behaviorally relevant stimuli that trigger fast alterations in gene expression patterns. Imaging of fluorescently tagged nucleosomes revealed that pharmacological manipulation of neuronal activity *in vitro* and auditory cued fear conditioning *in vivo* induce nucleus-scale restructuring of chromatin within minutes. Furthermore, the acquisition of auditory fear memory is impaired after infusion of a drug into auditory cortex which blocks chromatin reorganization *in vitro*. We propose that active chromatin movements at the nucleus scale act together with local gene-specific modifications to enable transcriptional adaptations at fast time scales. Introducing a transgenic mouse line for photolabeling of histones, we extend the realm of systems available for imaging of chromatin dynamics to living animals.

## Introduction

Interphase chromatin is organized in a highly ordered spatial structure with individual chromosomes occupying discrete chromosome territories within the nucleus (Bickmore and van Steensel, 2013; Cavalli and Misteli, 2013; Cremer and Cremer, 2001; Hubner and Spector, 2010). This three-dimensional nuclear architecture is believed to reflect the transcriptional state of the cell as the positioning of individual genes relative to transcriptional hotspots or the nuclear lamina is associated with their level of expression (Belyaeva et al., 2017; Buchwalter et al., 2019; Dekker and Mirny, 2016; Dekker and Misteli, 2015; Schoenfelder et al., 2010; Szabo et al., 2019; van Steensel and Belmont, 2017). Using microscopy, it has been demonstrated that the functional organization of interphase chromatin can undergo nucleus-scale plastic remodeling (Chuang et al., 2006; Csink and Henikoff, 1998; De Boni and Mintz, 1986; Kosak et al., 2002; Marshall et al., 1997; Mehta et al., 2010; Therizols et al., 2014; Wittmann et al., 2009) linked to changes in gene expression (Chambeyron and Bickmore, 2004; Chuang and Belmont, 2007; Chuang et al., 2006; Dion et al., 2012; Gasser, 2002; Gu et al., 2018; Martou and De Boni, 2000; Masui et al., 2011; Neumann et al., 2012; Thakar and Csink, 2005; Vazquez et al., 2001). Most of this work relied on the experimental accessibility of cultured cells.

It is well established, that changes in gene expression patterns in the brain are essential to support higher cognitive functions, such as the formation of long-term memories (Yap and Greenberg, 2018). Behavioral experiences, as well as specific patterns of neuronal activity, can induce changes in gene expression within few minutes (Abraham et al., 1993; Bertaina and Destrade, 1995; Guzowski et al., 1999; Peter et al., 2012). Furthermore, these changes are also reflected at the sub-microscopic level by alterations of the epigenetic landscape (Fischer, 2014; Halder et al., 2016; Yamada et al., 2019; Yang et al., 2016) and the conformation of chromosomes as described in recent studies using HiC techniques (Beagan et al., 2020; Marco et al., 2020; Watson and Tsai, 2017). However, it is not known, in how far fast, experience-induced changes in gene expression can be supported by sub-microscopic modifications at the molecular scale alone or are also reflected by and require nucleus-wide reorganization of chromatin as they have been described using microscopy in cultured cells. Currently, relatively little data on chromatin dynamics is available at the nucleus scale in tissue or even living animals (Barr and Bertram, 1949; Borden and Manuelidis, 1988). In particular, no longitudinal imaging of chromatin in individual neurons of living animals has been performed up to date.

In this study, we introduce a novel mouse model for photolabeling of chromatin to monitor the dynamics of chromatin in the mouse auditory cortex. We investigated in how far and at what time scales dynamic remodeling of chromatin can be observed using *in vivo* two-photon microscopy during basal conditions and auditory cued fear conditioning. We demonstrate that the rapid changes in gene expression patterns observed during memory formation are associated with nucleus-scale reorganization of chromatin structure in the brain.

## Results

### Nucleus-scale chromatin dynamics in primary cortical neurons

To investigate and visualize nucleus-scale chromatin dynamics in neurons upon induction of neuronal activity, we followed a widely used strategy that preserves endogenous nucleosome organization (Ricci et al., 2015): We transduced cultured primary cortical neurons (Blanquie et al., 2016) with a recombinant adeno-associated virus (rAAV) encoding for the core nucleosome component histone H2B fused to the fluorescent marker mCherry (H2B::mCherry) (Fig. 1A), and imaged labeled chromatin with a confocal microscope at a ten minutes interval. When cultures were kept in standard media, H2B::mCherry remained uniformly distributed in the nucleus except for regions within the nucleus that contain no or only little chromatin, e.g., the nucleolus (Bohn et al., 2010) (Fig. 1B top). When adding a pharmacological cocktail activating major neuromodulatory receptors and increasing synchronized network activity through a bicuculline-mediated block of inhibitory GABA_A_-receptors (Gutnick et al., 1982) (Fig. S1), we observed a fast increase in granularity of the fluorescent signal (Fig. 1B bottom, movie S1). Interestingly, this effect appeared to be transient as the chromatin label became more homogeneous again within the two-hour observation period.

**Fig. 1:**
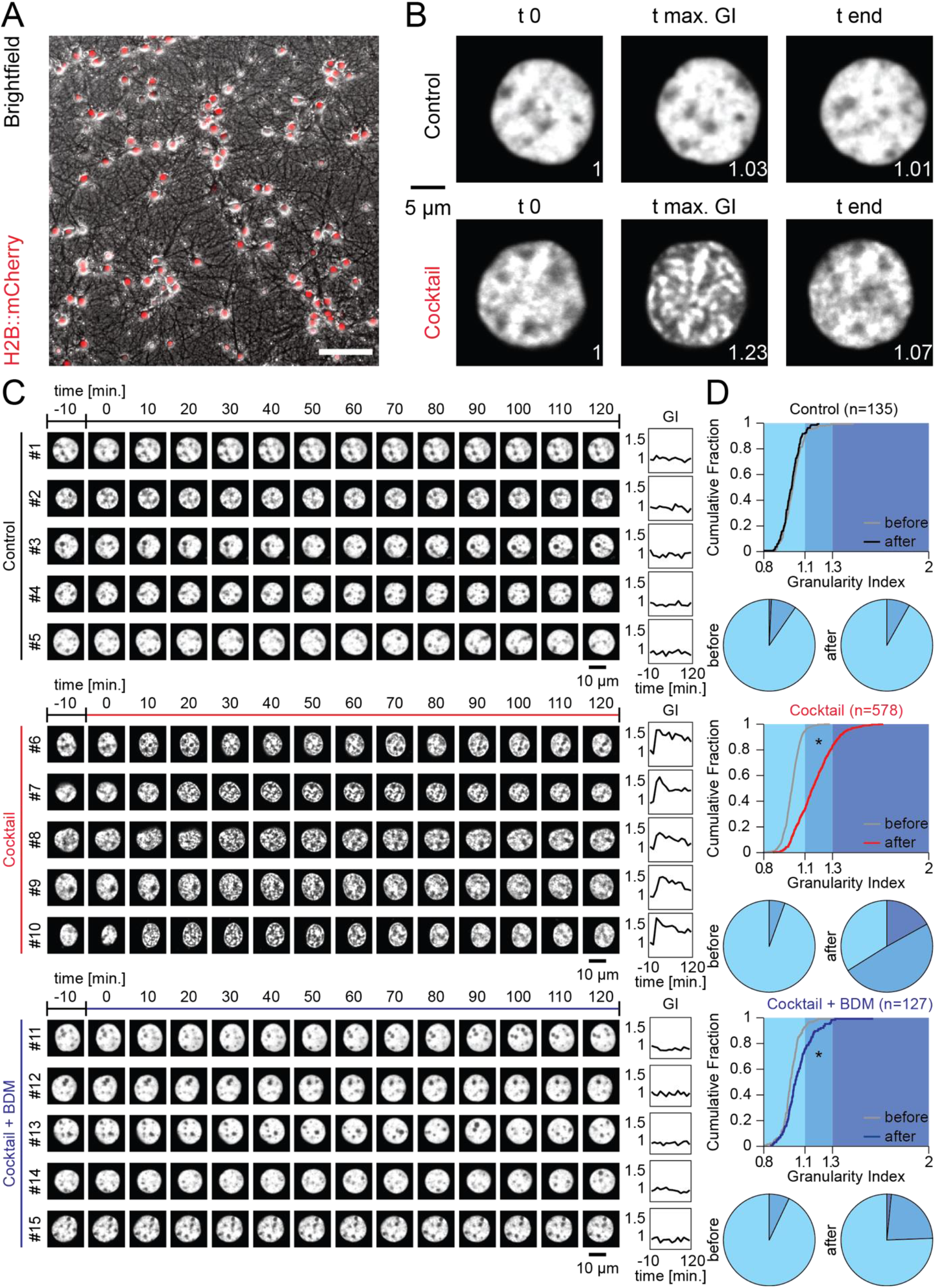
Transient reorganization of chromatin structure upon pharmacological induction of neuronal activity in primary cortical neurons. (**A**) Culture of primary cortical neurons expressing H2B::mCherry. Scale: 100μm. (**B**) Confocal images of two nuclei labeled with H2B::mCherry during a time lapse experiment. Top: Incubation with standard medium. Bottom: Incubation with pharmacological cocktail. First time point (t 0), time point of maximal Granularity Index (t max. GI), and last time point (t end). The corresponding GI value normalized to t 0 is shown on the bottom right of each image. (**C**) Time lapse images of example nuclei labeled with H2B::mCherry (Treatment conditions: control, black; cocktail, red; cocktail+BDM, blue). Colored line on top indicates treatment period. Right: time course of corresponding GI values normalized to t 0. (**D**) Cumulative distributions of GI values normalized to t 0 from all cells. Curves depict normalized GI values before (t −10; gray) and after application (t 40; control: black; cocktail: red; cocktail+BDM: blue). Bottom: Corresponding pie chart displays.

In order to quantify these changes in the nuclear distribution of H2B::mCherry, we calculated a Granularity Index (GI) based on the 2D Fourier transform of the fluorescent signal (for details see Methods) with larger GI values describing an increasingly uneven, granular distribution. GI values were normalized to time-point t 0, corresponding to the addition of the cocktail. When quantifying GI values in groups of neurons under cocktail conditions, we observed a systematic and significant increase in the GI, which peaked for most cells 40 minutes after application (Fig. 1, C, and D, movie S2). This increase was completely absent in cells under control conditions. We furthermore tested if the observed changes in chromatin condensation are dependent on actomyosin activity and included a third condition in which we applied the pharmacological cocktail together with BDM. This cell-permeable drug impairs myosin ATPase activity and has been shown to interfere with nuclear actomyosin dynamics underlying the movement of nuclear bodies (Chang et al., 2011; Fomproix and Percipalle, 2004; Muratani et al., 2002; Yarrow et al., 2003; Ye et al., 2008). The simultaneous application of the pharmacological cocktail with BDM significantly inhibited the activity-induced restructuring of chromatin (Fig. 1, C, and D) (control, n = 135: before: 1.017±0.007, after: 1.001±0.006; cocktail, n = 578: before: 1.005±0.003, after: 1.168±0.006; cocktail+BDM, n = 127: before: 1.000±0.006, after: 1.050±0.010; two-way ANOVA: factor time, d.f.: 1, F = 113.97, *p*<0.001, factor treatment, d.f.:2, F=88.88, *p*<0.001, interaction, d.f.: 2, F=104.11, *p*<0.001; differences between groups: control vs cocktail: before: *p*=0.969, after: *p*<0.001; control vs. BDM: before: *p*=1, after: *p*<0.001; cocktail vs. BDM: before: *p*=1, after: *p*<0.001; post-hoc multiple comparison test using Dunn and Sidák’s approach).

### Chromatin reorganization correlates with induction of c-Fos and increased γH2AX labeling

Next, we tested if activity-induced restructuring of chromatin is correlated with transcriptional activation. We repeated the previous experiments with the same conditions as described above (medium only, medium with the pharmacological cocktail, and medium with the pharmacological cocktail and BDM) and found a strong increase in the expression of the immediate-early gene product c-Fos after incubation with the pharmacological cocktail. The induction of c-Fos was blocked by the addition of BDM in a dose-dependent manner (Fig. 2, A, B, C, and D, S1, E, and F) (control, n = 90: 0.909± 0.051; cocktail, n = 372: 6.556± 0.251; cocktail+BDM, n = 106: 1.831± 0.202; ANOVA: d.f.: 2, F = 110.85, *p*<0.001, differences between groups: control vs. cocktail: *p*<0.001; control vs. cocktail+BDM: *p*=0.291; cocktail vs. cocktail+BDM: *p*<0.001; post-hoc multiple comparison test using Dunn and Sidák’s approach). Furthermore, the level of c-Fos immunoreactivity significantly correlated with the maximal GI of a given cell (Fig. 2D; n=485, R^2^=0.151, *p*<0.001).

**Fig. 2:**
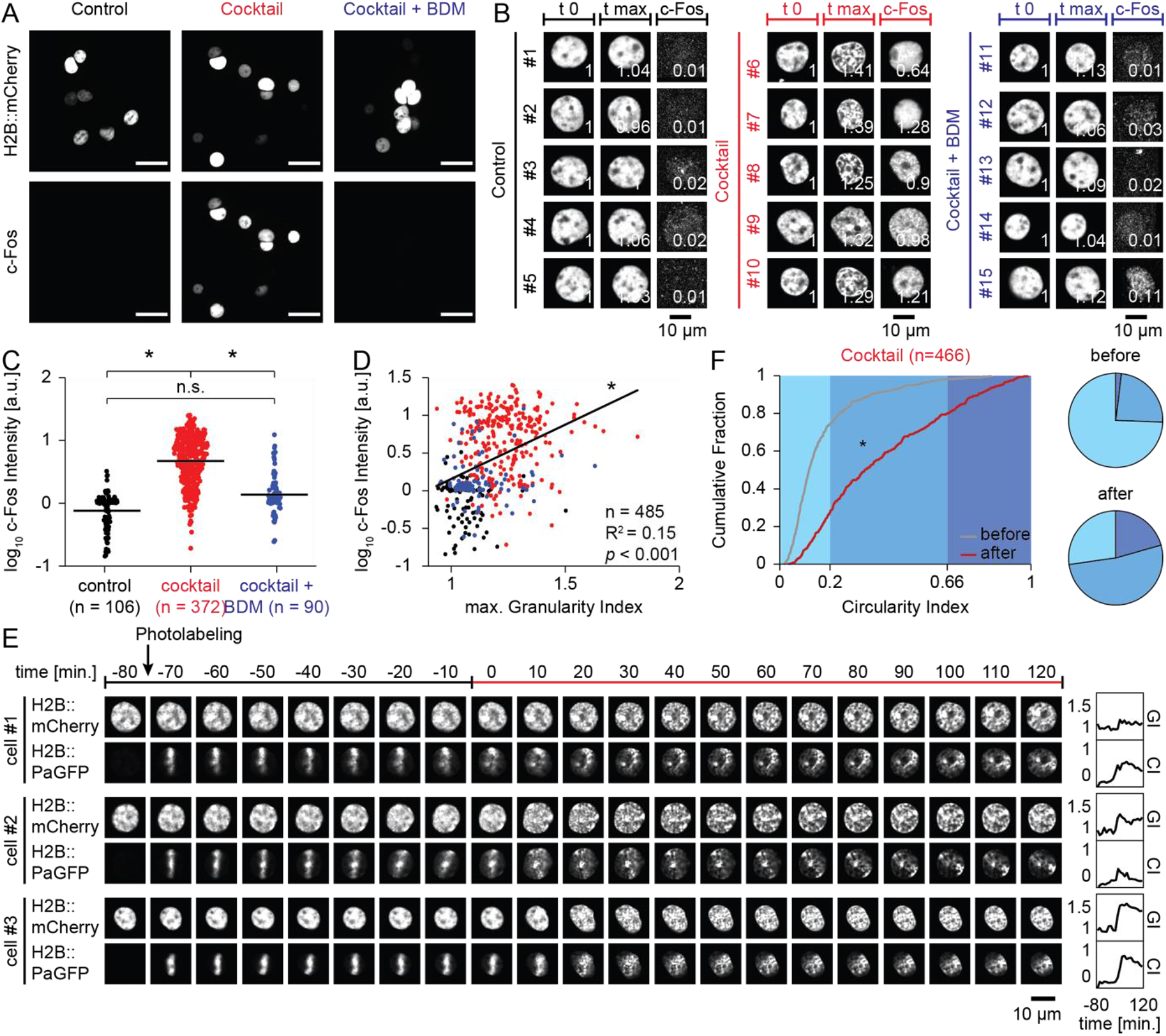
Reorganization of chromatin correlates with c-Fos induction and can be visualized by photolabeling of a fiducial marker. (**A**) Epifluorescence images of cultured primary cortical neurons expressing H2B::mCherry following treatment (control, cocktail, cocktail+BDM) and post-hoc immunofluorescence staining for the IEG product c-Fos. Scale bar: 25 μm. (**B**) Confocal time-lapse imaging of the H2B::mCherry signal and post-hoc c-Fos staining for example nuclei from the three treatment conditions (control, black; cocktail, red; cocktail+BDM, blue). Shown is H2B::mCherry signal at first time point (t 0) and time point of maximal normalized GI value (t max) during time-lapse imaging, as well as c-Fos staining after fixation. Bottom right of each image: Quantification of GI, or log_10_ c-Fos intensity, respectively. (**C**) Dot plot of log_10_ c-Fos intensity of all individual cells from the three treatment conditions (control, black; cocktail, red; cocktail+BDM, blue; horizontal lines indicate mean). (**D**) Scatter plot of maximal normalized GI value during time-lapse imaging and corresponding post-hoc log_10_ c-Fos intensity of individual cells. (control, black; cocktail, red; cocktail+BDM, blue; black line: linear fit). (**E**) Confocal time-lapse images of example nuclei co-expressing H2B::mCherry and H2B::PaGFP before and after incubation with pharmacological cocktail inducing neuronal activity. Top: arrow indicates photolabeling time point, colored line indicates treatment period. Right: Corresponding quantification of normalized GI and CI values over time. (**F**) Cumulative distribution of CI values from all cells. Curves depict CI value of each cell before (t −10; gray) and maximal CI value after time point of drug application (t 0 – t 120; red). Bottom: Corresponding pie chart displays.

Recent studies have shown increased occurrence of DNA double-strand breaks following neuronal stimulation and behavioral experience (Madabhushi et al., 2015; Suberbielle et al., 2013), a phenomenon classically associated to pathological conditions. Indeed, we observed an increase of γH2AX, a marker for DNA double-strand breaks, after induction of neuronal activity with the pharmacological cocktail. Consistent with our previous observations on c-Fos induction, the addition of BDM also inhibited the increase of γH2AX (Fig. S2).

### H2B::PaGFP facilitates tracking of chromatin reorganization

We wondered if nucleus-scale rearrangements of chromatin would be also triggered in the living brain by behaviorally relevant stimuli that are associated with changes in gene expression. The detection of chromatin reorganization using H2B::mCherry as marker requires imaging techniques with high spatial resolution restricting its applicability for *in vivo* approaches, where imaging conditions are typically less favorable compared to cell culture. To overcome this limitation, we followed a strategy to visualize chromatin dynamics by fusing H2B with a photoactivatable version of GFP (PaGFP) (Patterson and Lippincott-Schwartz, 2002). This approach leads to the integration of PaGFP into nucleosomes and allows for conditional photolabeling of fiducial makers to facilitate tracking of chromatin dynamics (Kanda et al., 1998; Strickfaden et al., 2010; Wiesmeijer et al., 2008).

We produced a rAAV encoding for the H2B::PaGFP fusion protein under the human Synapsin1 promoter, transduced primary cortical cultures with both the H2B::PaGFP and the H2B::mCherry viruses and performed similar time-lapse experiments as previously described. After the first imaging time point, we photolabeled thin lines across the nuclei of individual cells. We reasoned that the nucleus-scale reorganization of chromatin structure would be robustly detected as transformations of the shape of the photolabeled line. Similar to our previous observations, we only detected minimal changes in GI values in the H2B::mCherry channel during one hour of baseline imaging in standard medium and the labeled line in the H2B::PaGFP channel remained largely stable (Fig. 2E, movie S3). After addition of the pharmacological cocktail, we again observed an increase in granularity in the H2B::mCherry signal, which was paralleled by a drastic transformation of the photolabeled lines of H2B::PaGFP. In order to quantify changes of the photolabel, we derived a second metric, termed circularity index (CI). This metric is calculated as the ratio of the second to the first eigenvalue of a principal component analysis of the intensity distribution of the photolabel. This index is 0 for an infinitely thin, straight line and 1 for a circle. CI values began to rise sharply immediately after the application of the pharmacological cocktail, similar to the GI values (Fig. 2F; n = 466: before: 0.170±0.007, after: 0. 403±0. 012; ANOVA: d.f.: 1, F = 288.23, *p*<0.001, differences between groups: before vs. after: *p*<0.001; post-hoc multiple comparison test using Dunn and Sidák’s approach). This demonstrates that the transformation of the photolabeled fiducial marker captures a related phenotype of the same underlying process we had previously observed with increases in GI in the pan-nucleosome labeling approach using H2B::mCherry. In summary, using H2B::PaGFP for photolabeling of a fiducial marker and subsequent tracking of the transformation of this label allows robust detection of chromatin dynamics in neurons that is less dependent on optical resolution.

### Generation of a transgenic mouse line for photolabeling of chromatin

Having established the robust detection of chromatin dynamics using the CI of the H2B::PaGFP signal, we generated a conditional knock-in mouse line expressing H2B::PaGFP (Fig. S3 A, B, and C). The H2B::PaGFP expression pattern tightly correlated with the staining pattern of 4’,6-diamidino-2-phenylindole (DAPI), indicative of tight labeling of the chromatin structure in cortical neurons *in vivo*. After implantation of a cranial window, we successfully verified the selective, strong, and stable photoactivation of chromatin in cortical neurons (Fig. S3 D, and E). Furthermore, we did not detect changes in the electrophysiological properties of H2B::PaGFP expressing or photolabeled neurons (Fig. S4) and confirmed the ability to successfully acquire and express fear memories in mice that conditionally express H2B::PaGFP in excitatory neurons (Fig. S5).

### Activity-induced reorganization of chromatin in acute brain slices from transgenic mice expressing H2B::PaGFP

To compare the effects of neuronal activity on chromatin organization between dissociated neurons and the transgenic mouse model, we took advantage of an *in vitro* slice preparation, which allows measuring chromatin dynamics in isolated brain tissue. First, using paired patch-clamp whole-cell recordings in current-clamp mode, we observed that application of the pharmacological cocktail lead to reliable induction of synchronous bursting activity, similar to the previously described effects on dissociated cells (Fig. 3, A, B, C, and D; *p*<0.001, Fisher’s exact test). Furthermore, we observed that this treatment led to a robust induction of c-Fos immunoreactivity (Fig. S6). Additional bulk RNAseq in cocktail-treated and control slices showed differential expression of 205 genes (Fig. S7). We compared these results with a previous study in which the effects of various behavioral treatments on gene expression patterns in the auditory cortex *in vivo* were analyzed (Peter et al., 2012) and found that the application of the pharmacological cocktail leads to robust induction of many known immediate-early genes in brain slices similar to behavioral experiences *in vivo* (Fig. 3E).

**Fig. 3:**
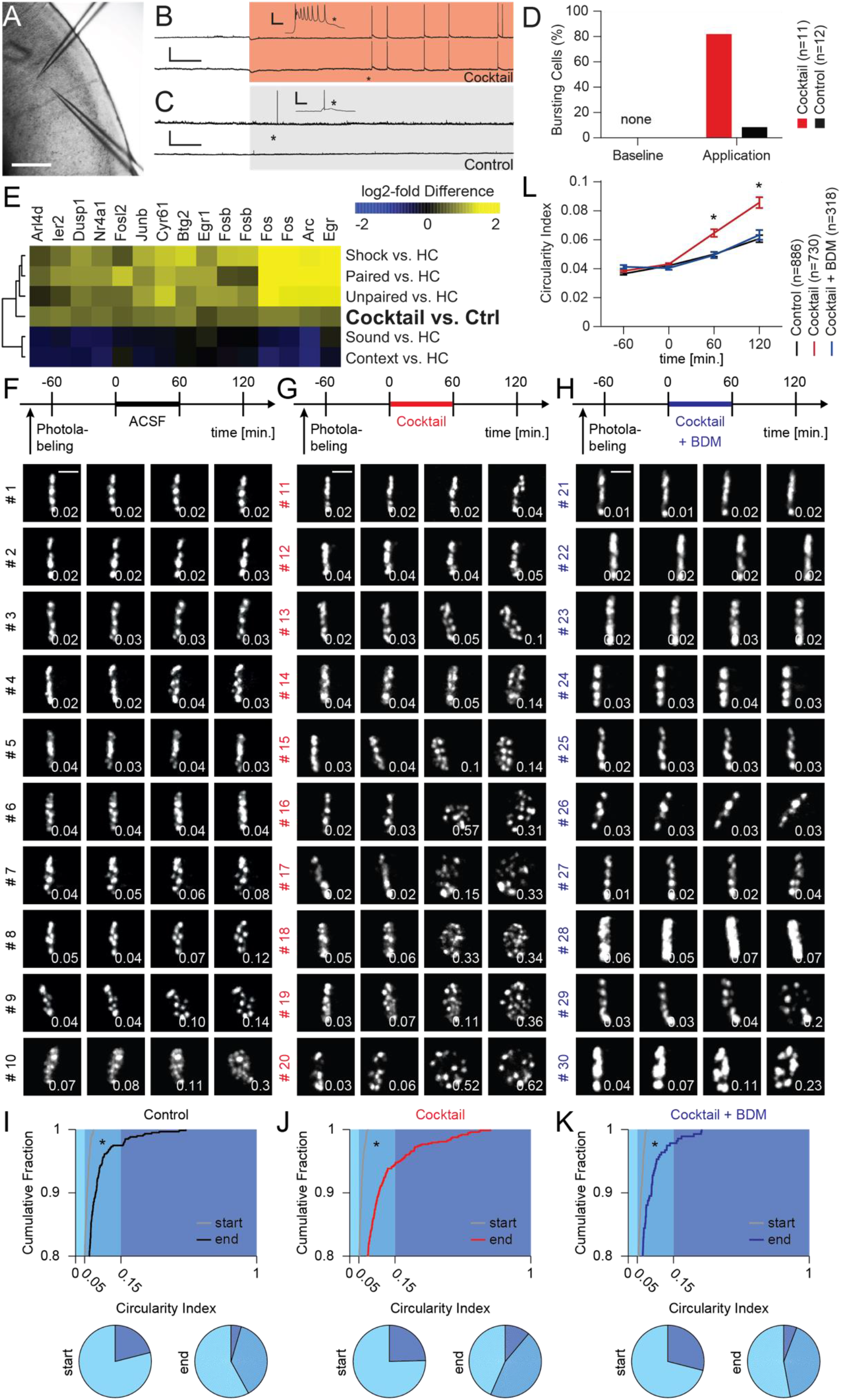
Pharmacological induction of neuronal activity in brain tissue induces chromatin reorganization. (**A**) Image of an acute brain slice of the auditory cortex from an H2B::PaGFP mouse with two patch pipettes illustrating recording configuration. Scale: 200μm. (**B**) Paired patch clamp whole-cell currentclamp recording of two neurons. Higher resolution of recordings indicated by asterisk. Red background indicates change of bath solution to ACSF containing a pharmacological cocktail driving synaptic transmission. Note, induction of highly synchronized busting activity during cocktail application. Scale: 40 mV, 2 min; scale inset: 30 mV, 100ms. (**C**) Analog control to the experiment shown in panel (b), however, normal ACSF was perfused during both baseline and application period. Note low activity levels in normal ACSF. (**D**) Quantification of burst events during baseline and application period in recordings from slices bathed with cocktail (red) and ACSF (black). (**E**) RNAseq following incubation (cocktail vs. control, six slices each) revealed a subcluster of significantly induced genes including many known IEGs. Additionally, corresponding relative expression levels of the genes are displayed that were observed in the auditory cortex following various behavioral treatments (data from (Peter et al., 2012)). Expression levels of some genes were measured with several probes on the microarray and are therefore mentioned multiple times. (**F**) Top panel: Timeline of the experiment to study chromatin dynamics in brain slices in the control group. Lower panels: examples of time-lapse of two-photon image stacks (maximum intensity projections) of individual photolabeled neurons, sorted by their respective CIs at 120 min. Scale bar: 5 μm. (**G**) Analog to panel (F), but for cells treated with pharmacological cocktail. (**H**) Analog to panel (F), but for cells treated with pharmacological cocktail+BDM. (**I**) Top: Cumulative distribution of all CIs at start (gray) and end (black) of the experiment from control slices. Bottom: Corresponding displays of the CIs at start and end of the experiment in pie chart format shown for the three groups. Category color code is indicated in plot above (light blue: < 0.05, medium blue: > 0.05 & < 0.15, dark blue: > 0.25). (**J**) Analog to panel (I), but for cells treated with pharmacological cocktail. (**K**), Analog to panel (I), but for cells treated with pharmacological cocktail+BDM. (**L**), Time course of mean CIs for control (black), cocktail (red) and cocktail+BDM (blue) treatment groups.

Next, we investigated the effect of increased neuronal activity on changes in chromatin structure. We labeled thin lines across nuclei of layers II/III neurons in the auditory cortex in acute brain slices obtained from H2B::PaGFP x CaMKIIa-Cre mice and acquired images at a one-hour interval for a total of four hours. Consistent with our findings in primary cortical neurons, we observed that the magnitude of chromatin reorganization as measured by changes in the CI was markedly different between control slices and slices treated with the pharmacological cocktail. Co-application of the cocktail together with BDM inhibited this effect (Fig. 3, F, G, H, I, J, K and L, S8; Control, n = 595: −60min, 0.037±0.001; 0min 0.042±0.001; 60min, 0.050±0.002; 120min, 0.061±0.003; Cocktail, n = 728; −60min, 0.039±0.001; 0min, 0.043±0.001; 60min, 0.065±0.003; 120min, 0.086±0.004; Cocktail + BDM, n = 276: −60min, 0.041±0.001; 0min, 0.040±0.001; 60min, 0.049±0.002; 120min, 0.063±0.003; two-way ANOVA: factor time, d.f.: 3, F = 103.8, *p*<0.001, factor treatment, d.f.:2, F=32.04, *p*<0.001, interaction, d.f.: 6, F=10.06, *p*<0.001; differences between groups: control vs cocktail: −60min: *p*=1, 0min: *p*=1, 60min: *p*<0.001, 120min: *p*<0.001; control vs. BDM: −60min: *p*=1,0min: *p*=1,60min: *p*=1, 120min: *p*=1; cocktail vs. BDM: −60min: *p*=1, 0min: *p*=1, 60min: *p*<0.002, 120min: *p*<0.001; post-hoc multiple comparison test using Dunn and Sidák’s approach). In contrast to dissociated cell cultures, in brain slices the treatment effects were more heterogeneous across neurons (Fig. S9): Whereas we also observed few cells acquiring high CI values even under control conditions, also many cells showed a stable photolabel following cocktail treatment. Together, these results demonstrate that patterns of neuronal activity that induce changes in gene expression induce at the same time nucleus-scale reorganization of chromatin in brain tissue.

### Dynamic chromatin organization *in vivo*

In order to investigate chromatin dynamics in a living animal, we injected rAAV-Cre into the auditory cortex of H2B::PaGFP mice, implanted a small cranial window above the auditory cortex (Loewenstein et al., 2011; Moczulska et al., 2013), and photolabeled the chromatin of cortical neurons with thin lines across the nucleus. We found that this strategy lead, in the vast majority of cases, to the labeling of neurons expressing the neuronal marker NeuN. (Fig. S10). When revisiting the same cells 24 hours after photolabeling, we observed that the line-pattern was essentially stable in many cells, while in others its shape was preserved as well, but rotated. In yet other cells, the photolabel was distributed across the nucleus, reminiscent of the observations in dissociated neurons and acute brain slices (Fig. 4A). A post-hoc analysis of *in vivo* photolabeled nuclei confirmed that an increase in the CI value was associated with a three-dimensional redistribution of the photolabel throughout the nucleus (Fig. S11). To obtain a better measure of the relevant time scales of chromatin reorganization *in vivo,* we monitored the transformation of the photolabel in an anesthetized mouse for three hours at a 15-minute interval (Fig. 4B). Similar to the findings in brain slices, chromatin dynamics were highly heterogeneous in neighboring neurons *in vivo*. In many cells, the label remained largely stable, while in others, the labeled line transformed and redistributed within the nucleus. When plotting the CI values of individual cells over time, we noted that phases of relative stability were interrupted by rapid and substantial transformations of the photolabel within few minutes (e.g. cells #5, #6 in Fig. 4B) (Chuang et al., 2006; De Boni and Mintz, 1986).

**Fig. 4:**
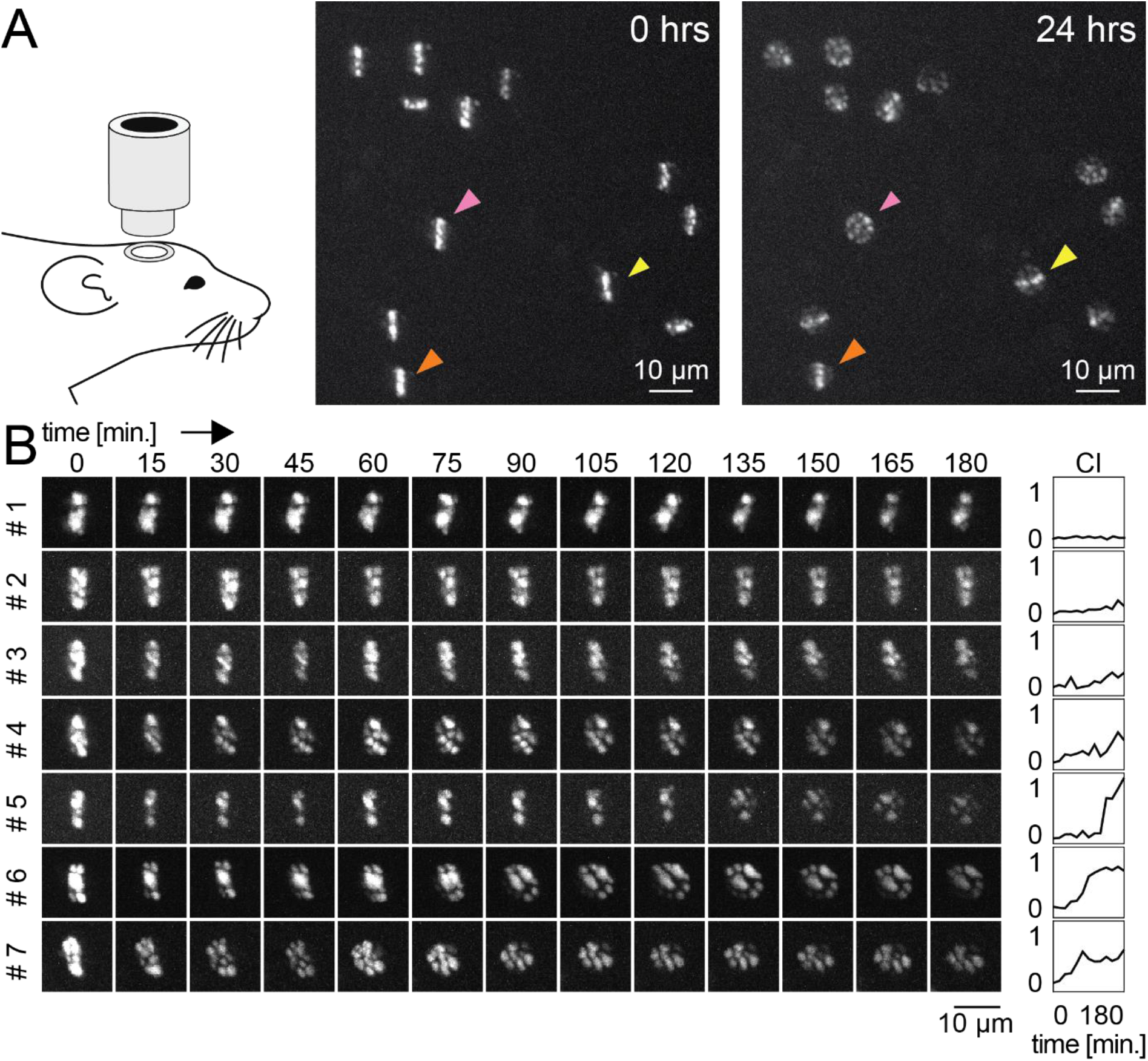
Nucleus-scale reorganization of chromatin *in vivo.* (**A**) Imaging of chromatin dynamics *in vivo* using a transgenic mouse line expressing H2B::PaGFP. Two-photon images of layer II/III auditory cortex neurons photolabeled with a line across the nucleus and imaged right after labeling (0 hrs) and a day later (24 hrs). Note substantial heterogeneity in the redistribution of labeled chromatin in individual neurons. Orange arrowhead: line pattern preserved, yellow arrowhead: line pattern preserved but rotated, pink arrowhead: uniform distribution of the line pattern. (**B**) Time-lapse maximum intensity projections of two-photon images taken *in vivo* from individual somata at a 15 min interval. Right panel: Corresponding CI quantification over time. CI remains stable over time or abruptly increases in individual neurons (e.g. #5-7).

Does the transformation of the shape of a photolabeled line *in vivo* reflect direct movements of chromatin or rather unbinding and re-binding of nucleosomes? In previous studies introducing photolabeling of tagged histones, only little diffusion of unbound nucleosomes has been reported and therefore, photolabeled fiducial markers have been used to track chromatin movements within the nucleus (Strickfaden et al., 2010; Wiesmeijer et al., 2008). Consistently, we could track individual speckles of photolabeled chromatin in our time-lapse data suggesting that changes in the CI reflect movements of chromatin rather than histone diffusion or turnover, which occurs on much slower time scales (Maze et al., 2015). Furthermore, the transformations of the patterns typically occurred in a directed manner, i.e. changes from image to image followed clear trajectories. This is in contrast to stochastic changes from image to image as they would be expected from a random walk-like diffusion process and rather suggests an actively driven process (Dion and Gasser, 2013; Marshall et al., 1997). We observed similar dynamics in wild-type mice using Hoechst to directly stain DNA *in vivo,* as an alternative method to track nucleus-scale movements of chromatin independent of nucleosome components tagged with a fluorophore (Fig S12). Passive diffusion of the photolabeled chromatin is expected to occur continuously in all cells. However, the observation that in some cells the photolabel can maintain a remarkable degree of stability over many hours indicates that, at the nucleus scale, only little chromatin motility results from passive processes alone. This suggests that under basal conditions chromatin appears to be highly viscous *in vivo* and effectively restrains diffusion (Chubb et al., 2002).

### Behavioral experiences induce nucleus-scale chromatin reorganization

To identify the physiological relevance of nucleus-scale chromatin dynamics, we combined *in vivo* imaging of chromatin with specific behavioral paradigms previously shown to induce plasticity in the mouse auditory cortex: Auditory cued fear conditioning and variants thereof affect the dynamics of synapses (Moczulska et al., 2013) and induce specific global patterns of gene expression (Fig. S13) (Peter et al., 2012). We stereotactically injected rAAV-Cre into the auditory cortex of H2B::PaGFP mice and implanted a cranial window. We used intrinsic imaging of sound-evoked activity patterns to map areas within the auditory cortex activated by the sound (CS+) used for conditioning (Fig. 5, A, B, and C) (Moczulska et al., 2013). We followed a longitudinal imaging strategy and exposed individual mice to a series of behavioral paradigms with increasing behavioral relevance (Fig. 5D). On each experimental day, we photolabeled thin lines on a set of cells in a small patch of layer II/III in the auditory cortex and acquired image stacks of those labeled cells directly after photolabeling and two hours later.

**Fig. 5:**
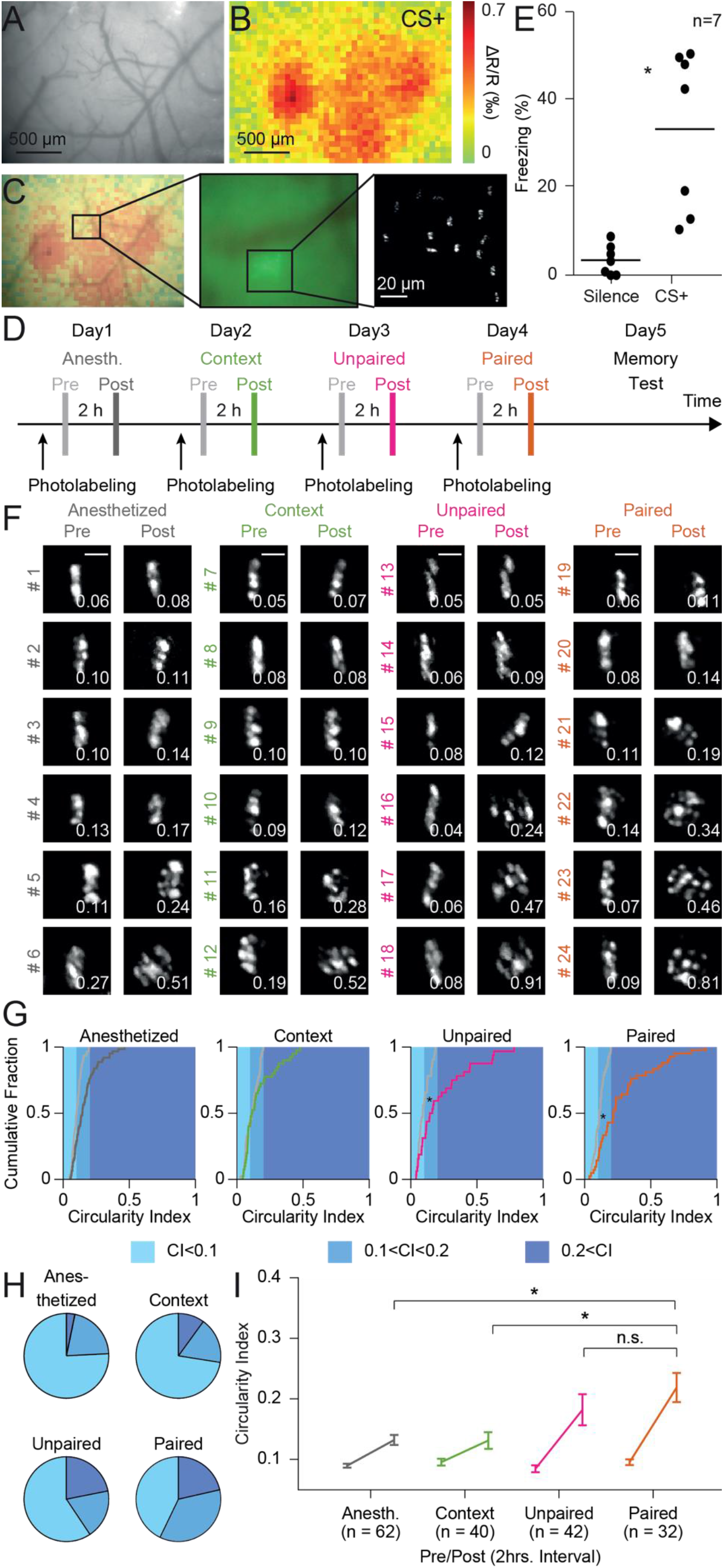
Induction of chromatin reorganization by behavioral experiences. (**A**) Pattern of surface vessels over the auditory cortex as seen through a cranial window. (**B**) Same field of view displaying intrinsic signals indicating metabolic activation by a sound used as conditioned stimulus (CS+) for auditory cued fear conditioning. (**C**) Left: Overlay of vessel pattern and intrinsic optical signals. Middle: Epifluorescence image with shadows of surface vessels and field of photolabeled neurons. Right: Labeled neurons in a two-photon image of layers II/III. (**D**) Schematic of the experimental timeline. Chromatin was imaged directly after (Pre) and two hours after (Post) photolabeling. Different behavioral treatments were performed between the two imaging sessions. (**E**) Quantification of freezing behavior of mice in a memory test session one day after photolabeling and auditory cued fear conditioning. (**F**) Maximum intensity projections of two-photon image stacks of individual photolabeled neurons directly after labeling (Pre) and 2 hours later (Post). Mice were either anesthetized, placed in a known context, subjected to unpaired auditory cued fear conditioning, or subjected to paired auditory cued fear conditioning, as indicated in the headings. Cells are sorted by CI values at the second time point. Scale bar: 5 μm. (**G**) Cumulative distribution plots of CIs of all neurons labeled for the four experimental conditions. Gray curves indicate CIs directly after photolabeling, the colored lines the CIs of the same cells after the 2-hour interval. (**H**) Corresponding displays of the CIs at the second time point in pie chart format. Color code is indicated in panel g (light blue: < 0.1, medium blue: > 0.1 & < 0.2, dark blue: > 0.2). (**I**) CIs of photolabeled neurons directly after labeling and after 2-hour interval. Data is from seven mice.

On the first day, animals were kept under the microscope under isoflurane anesthesia during the two hours as baseline control. On the second day, mice were woken up after the first imaging time point and returned to their home cage, a familiar environment. On the third day, experimental subjects underwent unpaired auditory cued fear conditioning between the two imaging time points, in which five sound presentations and foot shocks were applied temporally separated by more than a minute. Unpaired auditory cued fear conditioning does not lead to the specific formation of a tone-shock association but represents a stressful experience. It has previously been shown that unpaired auditory cued fear conditioning induces an extensive change of the transcriptional program (Peter et al., 2012), and increased rewiring of neuronal connections (Moczulska et al., 2013) in the auditory cortex. On the fourth day, mice underwent paired auditory cued fear conditioning between the two imaging time points, in which the conditioned sound was directly followed by a mild foot shock. Successful memory formation was assessed in a memory test session in a neutral environment on the next day. Here, we observed an increase in freezing behavior during the presentation of the conditioned sound, demonstrating the successful formation of an associative memory (Fig. 5E; Freezing levels: silence: 0.043±0.016, CS+: 0.416±0.086, n=7, ANOVA: d.f.: 1, F = 18.05, *p*<0.002, differences between groups: baseline vs. CS+: *p*<0.002). When analyzing the CIs of the line patterns, a two-way ANOVA revealed a significant effect of time (d.f.: 1, F=67.48, *p*<0.001), treatment (d.f.:3, F=6.09, *p*<0.001) and a significant interaction (d.f.: 3, F=5.74, *p*<0.001) (Fig. 5, F, G, H, and I).

Increases in the CIs were observed during the two hours imaging interval in all groups but did not reach statistical significance for cells of mice which were anesthetized or were put into the familiar context. (Fig. 5I; anesthetized, n=62, pre: 0.105±0.005, post: 0.164±0.012, *p*=0.122; context, n=40, pre: 0.113±0.008, post: 0.163±0.019, *p*=0.793; unpaired, n=32, pre: 0.098±0.008, post: 0.234±0.036, *p*<0.001; paired, n=42, pre: 0.113±0.007, post: 0.285±0.034, *p*<0.001; post-hoc multiple comparison test using Dunn and Sidák’s approach). The fraction of neurons showing high CI values was significantly larger with increased behavioral relevance of the paradigm. In the two-hour time interval, significantly more cells showed stronger increases when the mice underwent unpaired and paired conditioning, the two paradigms that lead to large changes in transcriptional activity, in comparison to the anesthetized and context groups (Fig. 5I; anesthetized post vs. unpaired post, *p*=0.155, anesthetized post vs. paired post, *p*<0.001, context post vs. unpaired post, *p*=0.252, context post vs. paired post, *p*<0.001, unpaired post vs. paired post, *p*=0.825; post-hoc multiple comparison test using Dunn and Sidák’s approach). We did not detect a significant difference between the unpaired and the paired group, the two paradigms with the highest behavioral relevance. These results show that not only increased neuronal activity in primary neurons and brain slices, but also behaviorally relevant experiences that induce plasticity in synapses and gene expression profiles control the rate of nucleus-scale chromatin dynamics *in vivo*.

### Inhibition of chromatin dynamics in auditory cortex impairs memory formation but not memory retrieval

Changes in the transcriptional program are necessary for the acquisition and consolidation of long-term memories (Yap and Greenberg, 2018). Based on our findings that the inhibitor of nuclear actomyosin dynamics BDM efficiently blocks activity-induced chromatin reorganization and c-Fos expression, we hypothesized that interference with chromatin dynamics could impair memory acquisition. To test this, we bilaterally implanted cannulas on top of the auditory cortex of wild-type mice (Fig. S14) and subjected them to an auditory cued fear conditioning task. Immediately before the first conditioning session, we either infused BDM to block chromatin reorganization or with ACSF as control and probed the memory on the next day (Fig. 6A). Consistent with our hypothesis, freezing levels were significantly lower in the BDM group during CS presentation but not baseline, indicating an impairment of fear memory (Fig. 6B left). To test if this effect reflects a specific deficit in memory formation and not an impairment of memory maintenance or sound processing necessary for cued fear expression (Heale and Harley, 1990), we subjected the same cohort of mice to a second conditioning session without pharmacological treatment. Prior to the memory test session on the next day, we infused the auditory cortex with either BDM or ACSF and subjected the two groups to the conditioned stimulus to assess memory retrieval. While Group 2 recovered from the memory deficit, we did not detect significant differences in freezing levels within between the BDM and ACSF treated groups. These data indicate that BDM does not interfere with memory retrieval or a behavioral fear response (Fig. 6B right; freezing levels: memory test 1, baseline: group 1, n=12, 10.167± 3.548, group 2, n=10, 7.400±3.444; CS: group 1,56.060±7.002, group 2, 34.963±2.882; memory test 2, baseline: group 1, 9.435±2.529, group 2, 5.042±2.041; CS: group 1, 56.188±7.316, group 2, 44.656±3.748; RM three-way ANOVA: period: d.f.: 1,20 F = 171.5, *p*<0.001, session: d.f.: 1,20 F = 0.235, *p*=0.633, group: d.f.: 1,20 F = 0.351, *p*=0.56, period vs. session: d.f.: 1,20 F=1.287, *p*=0.270, period vs. group: d.f.: 1,20 F=0.841, *p*=0.370, session vs. group: d.f.: 1,20 F=8.21, *p*=0.010, period vs. session vs. group: d.f.: 1,20 F=5.01, *p*=0.037; differences between groups: memory test 1 baseline, group 1 vs group 2: *p*>0.999, memory test 2 baseline, group 1 vs. group 2: *p*>0.999; memory test 1 CS, group 1 vs group 2: *p*=0.019, memory test 2 CS, group 1 vs. group 2: *p*=0.611; differences within groups: memory test 1, group 1, CS vs. memory test 2, group 1, CS: *p*=0.564; memory test 1, group 2, CS vs. memory test 2, group 2, CS: *p*=0.032; post-hoc multiple comparison test using Sidák’s approach). Taken together, inhibition of chromatin dynamics in vivo significantly impairs memory formation but not memory retrieval in mice.

**Fig. 6:**
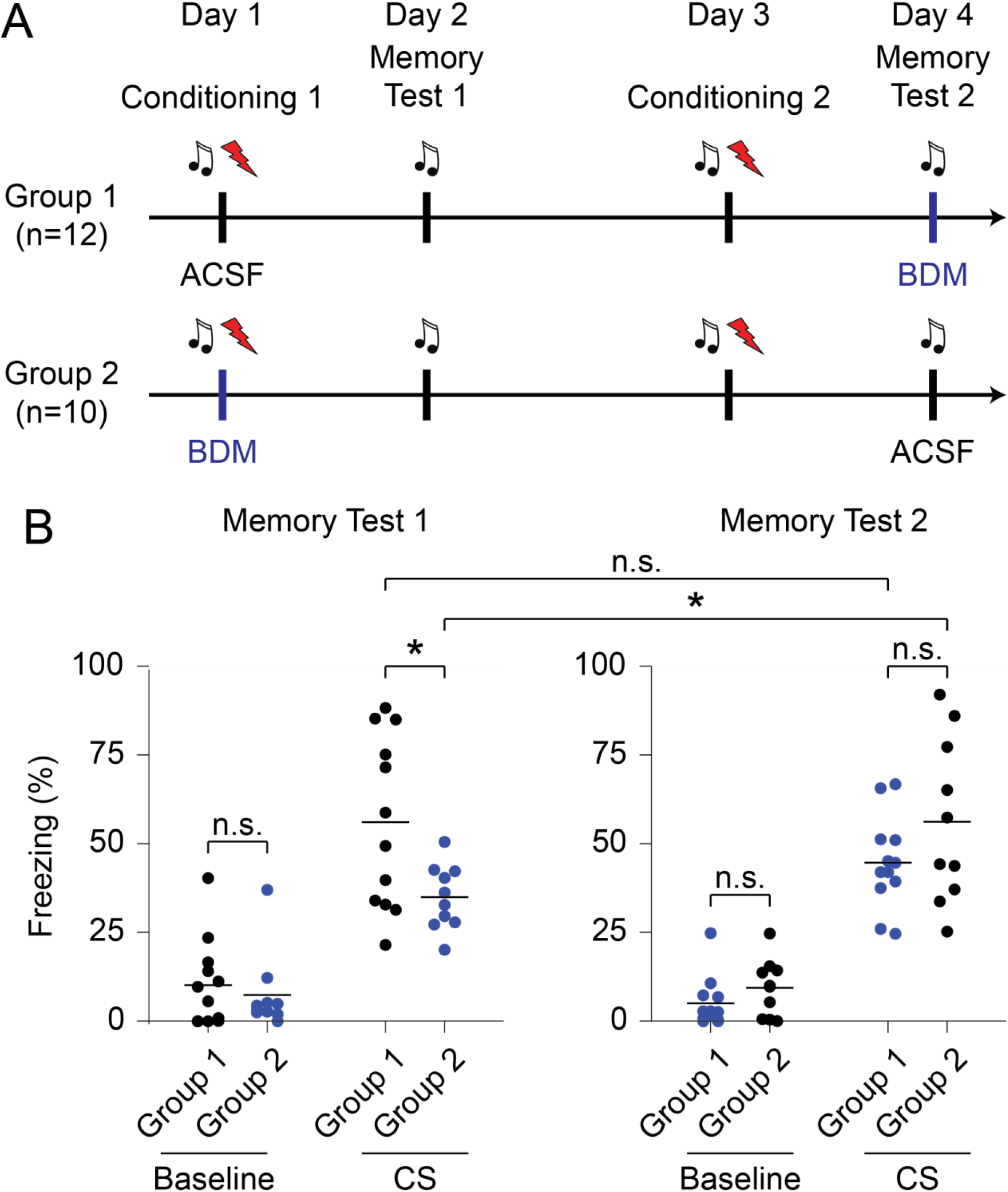
BDM mediated inhibition of chromatin dynamics in auditory cortex impairs memory formation, but not retrieval. (**A**) Schematic of experiment: Group 1 was infused with ACSF during first conditioning session, and with BDM during second memory test. Group 2 was infused with BDM during first conditioning session, and with ACSF during second memory test. Each behavioral session was separated by one day. (**B**) Quantification of freezing behavior of mice during both memory tests. Shown is freezing during first 30 seconds baseline in silence and during replay of the conditioned sound stimulus. Black: ACSF; blue: BDM.

## Discussion

Here, we show that changes in patterns of neuronal activity, as well as specific behavioral experiences, are correlated with a rapid, nucleus-scale reorganization of chromatin as well as changes in the transcriptional program of the cells. Furthermore, BDM, an inhibitor of nuclear forms of myosin, blocks chromatin dynamics, prevents the induction of the IEG c-Fos, and impairs the formation of long-term memories, whose consolidation is known to depend on transcriptional regulation. Together, this study demonstrates the requirement of nucleus-scale chromatin rearrangements at the microscopic level to allow transcriptional changes underlying experience-dependent behavioral adaptation.

To extend the systems available for the study of chromatin dynamics from cultured cells to tissue preparations and living animals, we generated a novel transgenic mouse line that allows conditional expression of the photoactivatable fluorescent protein, PaGFP, fused to the histone H2B. Apart from the analysis of chromatin per se, this mouse line offers substantially enhanced labeling of individual cells for various photolabeling applications where reliable tracking and re-identification of cells is crucial (Peter et al., 2013).

How could nucleus-scale reorganization of chromatin contribute to alterations in the transcriptional program necessary for behavioral adaptation? There are three scales to consider. First, the molecular scale of individual genes: There is good evidence that the transcriptional activity in neurons, similar to other cell types, is controlled by epigenetic modifications and changes in affinities of promoters and enhancers to transcription factors (Fischer, 2014; Korzus et al., 2004; Levenson et al., 2004; Miller et al., 2010; Peleg et al., 2010; Riccio, 2010; Vialou et al., 2013). Second, the nucleus scale: The localization of individual genes into functionally active domains, such as euchromatin near nuclear speckles, or functionally repressed domains, such as heterochromatin close to the lamina, occurs at the micrometer scale (Bickmore and van Steensel, 2013; Cavalli and Misteli, 2013; Cremer and Cremer, 2001; Dion and Gasser, 2013; Hubner and Spector, 2010; Lanctot et al., 2007). Third, the temporal scale: The activation of IEGs (Abraham et al., 1993; Bertaina and Destrade, 1995; Guzowski et al., 1999; Peter et al., 2012) and corresponding chromosome conformational changes (Beagan et al., 2020) are reported within minutes after induction. Our observation that the chromatin architecture at the nucleus scale appears to be little affected by passive diffusion-like processes (Fig. 4A, orange arrowhead) but can be rapidly adapted in response to behavioral experience, connects these three scales in the living animal.

Our findings are consistent with a model in which neuronal activity triggers processes dependent on nuclear myosin that put the nucleus in a permissive state allowing local, gene specific modifications of promoters and other regulatory elements to fall into a new organizational structure of the nucleus, resulting in an adaptation of the transcriptional program. Interestingly, nuclear forms of myosin have been recently implicated as autism spectrum disorder risk genes (Yuen et al., 2017). In addition to myosin-dependent processes, recent studies implicate a phase separation mechanism to create such a permissive state (Banani et al.; Boija et al., 2018; Cho et al., 2018; Gibson et al., 2019; Hnisz et al., 2017; Larson et al., 2017; Sabari et al., 2018; Sanulli et al., 2019; Shrinivas et al., 2019; Strom et al., 2017). Furthermore, our findings relate to the report of increased occurrence of double-strand breaks in neurons of mice exposed to a novel environment (Suberbielle et al., 2013): Global nucleus-scale movements likely lead to increased chromatin tension causing topological constraints for transcription of activated genes which would require resolution by double-strand breaks (Madabhushi et al., 2015). In recent years, HiC studies have allowed detailed insight into molecular features of chromatin reorganization upon transcriptional activation (Marco et al., 2020). However, as these techniques typically rely on pooling of cells and fixating them at a given time point, our *in vivo* imaging approach provides a valuable complement enabling the longitudinal study of chromatin dynamics in individual cells and thereby to explore their heterogeneity within a tissue.

In summary, we report rapid nucleus-scale chromatin dynamics in a living animal, which are associated with fast changes in transcriptional states and are essential for successful behavioral adaptation. Although, we focused in our study on memory formation, we believe that nucleus-scale reorganization of chromatin could be a general process associated to fast changes in the transcriptional program also in other contexts.

## Author Contributions

MP designed the experiments, characterized H2B::PaGFP in cell culture, produced and characterized the transgenic mouse model, performed *in vivo* chromatin labeling experiments, behavioral experiments, NeuN immunohistochemistry, confocal imaging, analyzed data, and prepared the manuscript; DA designed the experiments, analyzed chromatin labeling experiments in primary cultures, performed and analyzed chromatin labeling experiments in brain slices as well as *in vivo,* behavioral experiments, confocal imaging, cannula histology, analyzed data, and prepared the manuscript; RVP performed chromatin labeling experiments in primary cultures; AS performed and supervised experiments in primary cultures; FG performed brain slice electrophysiology and c-Fos immunohistochemistry; DK performed and analyzed behavioral experiments with cannulas; KK performed behavioral experiments; TB analyzed RNAseq data; HJL supervised chromatin labeling experiments in primary cultures and edited the manuscript; WH supervised behavioral experiments the immunohistochemistry and RNAseq experiments, and edited the manuscript; SR designed the experiments, was responsible for the overall project strategy and management, and prepared the manuscript.

## Acknowledgements

The authors would like to thank Isabel Bauer, Simone Dahms-Praetorius, Beate Krumm, Violetta Steinbrecher, Barbara Werner and Andreas Sommer for help with experiments, Christof Cremer, Thomas Cremer and Kurt Kremer for helpful discussions and Helmut Kessels for careful reading of an earlier version of the manuscript. WH and FG were supported by the European Community’s Seventh Framework Programme (FP/2007-2013) / ERC grant agreement no. 311701. MP and SR were supported by the Austrian Science Fund (FWF): P21930-B09. AS, HJL and SR were supported by the Deutsche Forschungsgemeinschaft (DFG) collaborative research center CRC1080 (TP A01 and TP C05). Support by the Core Facility Microscopy of the IMB in Mainz is gratefully acknowledged. RVP is a graduate student of the International PhD programme coordinated by the Institute of Molecular Biology Mainz. The Research Institute of Molecular Pathology (IMP) is supported by Boehringer Ingelheim and the Austrian Research Promotion Agency (FFG). The Vienna BioCenter Core Facilities (VBCF) Preclinical Phenotyping Facility acknowledges funding from the Austrian Federal Ministry of Education, Science & Research; and the City of Vienna.

## Supplementary Information

**Movie S1: Time-lapse video of the cells shown in Fig. 1B.**

**Movie S2: Time-lapse video of the cells shown in Fig. 1C.**

**Movie S3: Time-lapse video of the cells shown in Fig. 2E.**

**Fig. S1:**
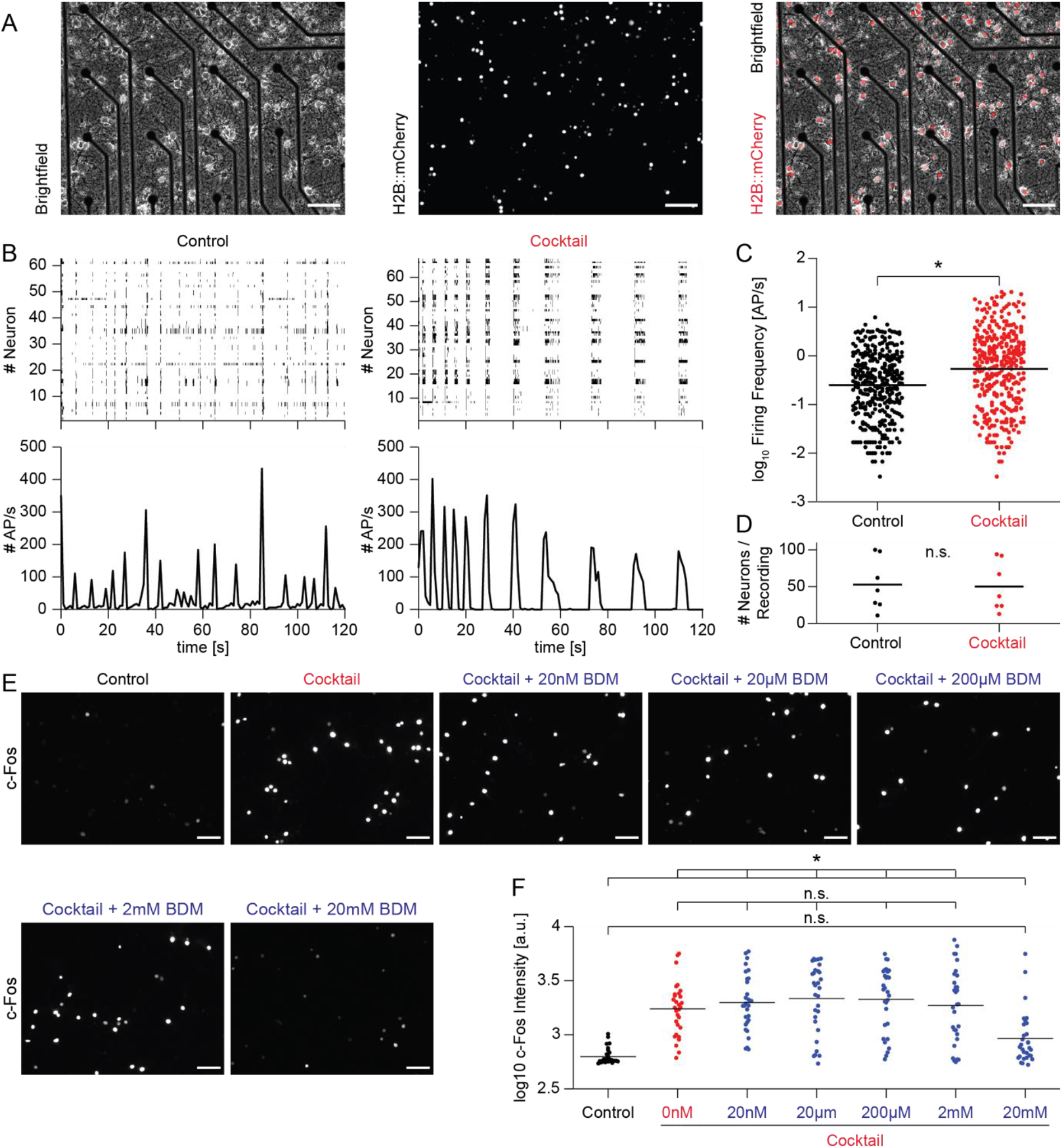
Pharmacological cocktail increases synchronized network activity in primary cortical cultures and c-Fos induction is inhibited by BDM in a dose-dependent manner. (**A**) Epifluorescence images of cultured primary neurons on a multielectrode array (MEA) with rAAV-mediated expression of H2B::mCherry. Left: Brightfield, middle: H2B::mCherry, right: merged image; scale bar: 50 μm. (**B**) Top: Raster plots of single unit firing rates recorded on a MEA array under two different conditions. Bottom: Summed number of action potentials from all recorded single units in experiment shown on top. Left: control, right: cocktail. (**C**) Quantification of single unit firing frequency from all seven recordings (control, n=370, mean±sem: 0.635±0.045; cocktail, n=351, mean±sem: 1.99±0.19). Firing frequency is significantly increased during cocktail incubation (ANOVA, factor group, d.f.: 1, F=36.63, *p*<0.001). (**D**) Comparable number of neurons per MEA recording (control, n=7 recordings, mean±sem: 52.86±13.36; cocktail, n=7 recordings, mean±sem: 50.14±12.80). The number of neurons per recording is not significantly different between groups (ANOVA, factor group, d.f.: 1, F=0.01, *p*=0.917). (**E**) Epifluorescence images of immunofluorescence staining against c-Fos after incubation of primary neurons with the pharmacological cocktail in combination with different concentrations of BDM. Respective condition is indicated on top of each image. (**F**) Quantification of c-Fos intensity for different conditions from panel (E) (control, n=30, mean±sem: 636.68±22.32; cocktail, n=30, mean±sem: 2027.02±230.43; cocktail+20nM, n=30, mean±sem: 2362.70±267.47; cocktail+20μM, n=30, mean±sem: 2692.40±282.05; cocktail+200μM, n=30, mean±sem: 2582.23±264.83; cocktail+200μM, n=30, mean±sem: 2497.58±351.20; cocktail+20mM, n=30, mean±sem: 1133.77±193.57). c-Fos expression is not significantly different between control and cocktail+20mM BDM. All other conditions show significantly increased c-Fos expression (ANOVA, factor group, d.f.: 6, F=18.41, *p*<0.001; differences between groups: control vs. cocktail: *p*<0.001; control vs. cocktail+20nM: *p*<0.001; control vs. cocktail+20μM: *p*<0.001; control vs. cocktail+200μM: *p*<0.001; control vs. cocktail+2mM: *p*<0.001; control vs. cocktail+20mM: *p*=0.299; cocktail vs. cocktail+20nM: *p*=1; cocktail vs. cocktail+20μM: *p*=0.975; cocktail vs. cocktail+200μM: *p*=0.992; cocktail vs. cocktail+2mM: *p*=1; cocktail vs. cocktail+20mM: *p*=0.002; cocktail+20nM vs. cocktail+20μM: *p*=1; cocktail+20nM vs. cocktail+200μM: *p*=1; cocktail+20nM vs. cocktail+2mM: *p*=1; cocktail+20nM vs. cocktail+20mM: *p*<0.001; cocktail+20μM vs. cocktail+200μM: *p*=1; cocktail+20μM vs. cocktail+2mM: *p*=1.0; cocktail+20μM vs. cocktail+20mM: *p*<0.001; cocktail+200μM vs. cocktail+2mM: *p*=1; cocktail+200μM vs. cocktail+20mM: *p*<0.001; cocktail+2mM vs. cocktail+20mM: *p*<0.001).

**Fig. S2:**
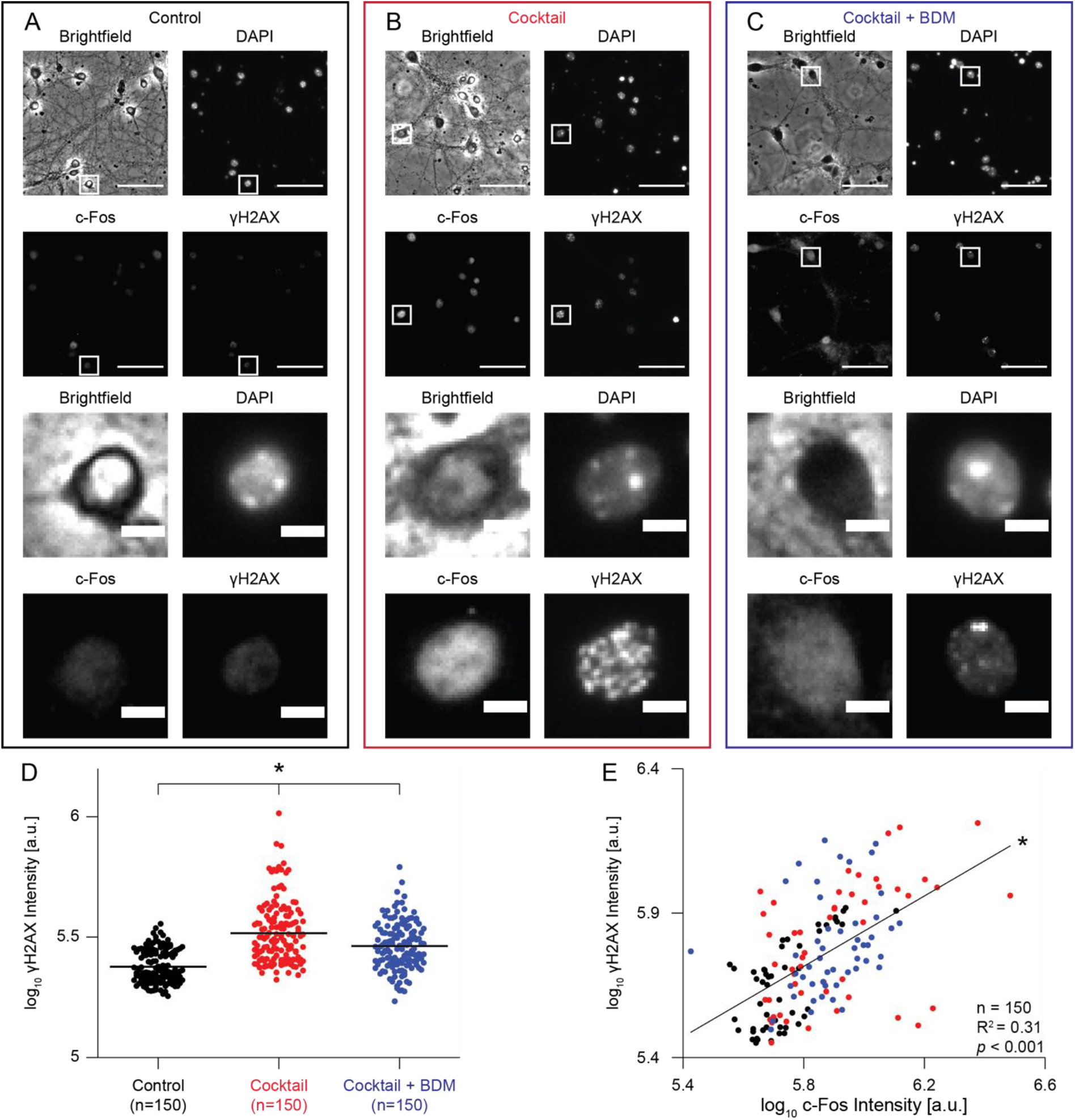
Neuronal activity induces γH2AX, a marker of DNA double-strand breaks. (**A**) Epifluorescence images of fixed cultured primary neurons from control condition after immunofluorescence staining for c-Fos and γH2AX, additionally stained with DAPI. Respective fluorescence channel is indicated on top of image. Scale bar: 50 μm. White squares indicate representative nucleus shown at higher magnification in bottom four images. Scale bar: 5 μm. (**B**) Same as panel (A) for cocktail condition. (**C**) Same as panel (A) for cocktail+BDM condition. (**D**) Dot plot of log_10_ γH2AX intensity for 150 individual cells from three different groups, respectively (control, black; cocktail, red; cocktail+BDM, blue; control: mean±sem: 301750±4991.2; cocktail: mean±sem: 477180±18360; cocktail+BDM: mean±sem: 394460±9659.3). γH2AX expression is significantly different between control and cocktail, control and cocktail+BDM, as well as between cocktail and cocktail+BDM (ANOVA, factor group, d.f.: 2, F=73.26, *p*<0.001; differences between groups: control vs. cocktail: *p*<0.001; control vs. cocktail+BDM: *p*<0.001; cocktail vs. cocktail+BDM: *p*<0.001). (**E**) Scatter plot of post-hoc log_10_ c-Fos intensity and log_10_ γH2AX intensity in 50 cells per group (n=150 cells; black line: linear fit, R^2^=0.31, *p*<0.001). Group color code is the same as in panel (D).

**Fig. S3:**
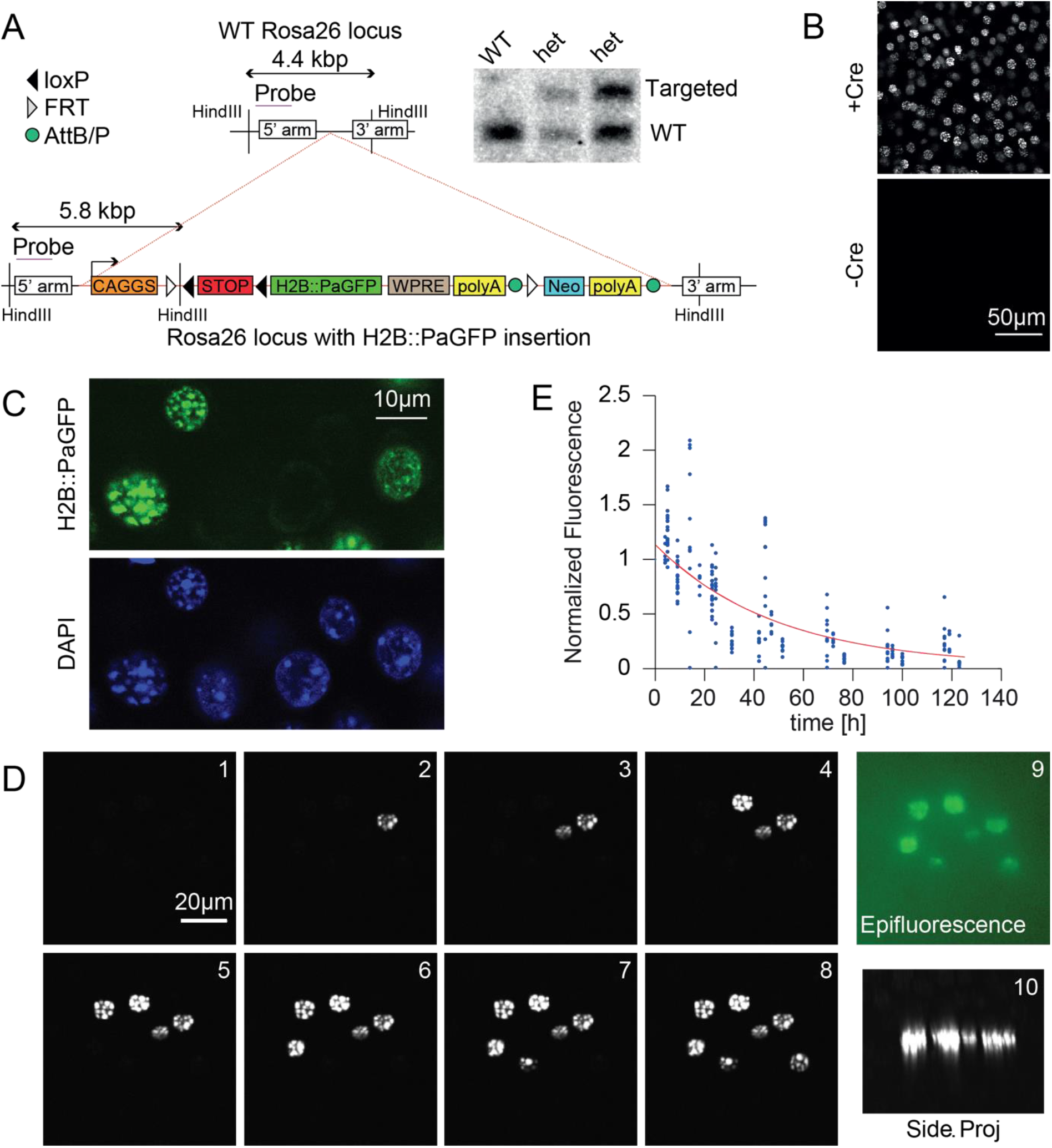
A transgenic mouse model to photolabel chromatin. (**A**) Targeting strategy for the Rosa26 locus and southern blots of wild-type and H2B::PaGFP heterozygous mice. H2B::PaGFP is expressed from the CAGGS promoter, directly upstream of the transgene is a floxed STOP cassette. (**B**) Top panel: Confocal image of a fixed section of the auditory cortex from a H2B::PaGFP mouse crossed with an EMX1-Cre mouse line after broad photolabeling. Bottom panel: Analog image from a Cre negative H2B::PaGFP littermate. (**C**) Confocal image of a fixed section of the auditory cortex taken from a H2B::PaGFP mouse with rAAV-mediated Cre expression. Top panel: H2B::PaGFP fluorescence after broad photolabeling in the imaging plane, bottom panel shows DAPI fluorescence. A strong correlation between both signals can be observed in transduced cells. (**D**) Panel 1-8: Maximum intensity projection from *in vivo* two-photon image stacks of consecutively photolabeled nuclei in the auditory cortex. Note selective activation of H2B::PaGFP fluorescence in nuclei which have been photoactivated at 750 nm. Panel 9: In vivo epifluorescence image of the same activated nuclei. Panel 10: Side projection of an image stack of the activated nuclei from panel 8, illustrating selective photoactivation in the imaging plane.

**Fig. S4:**
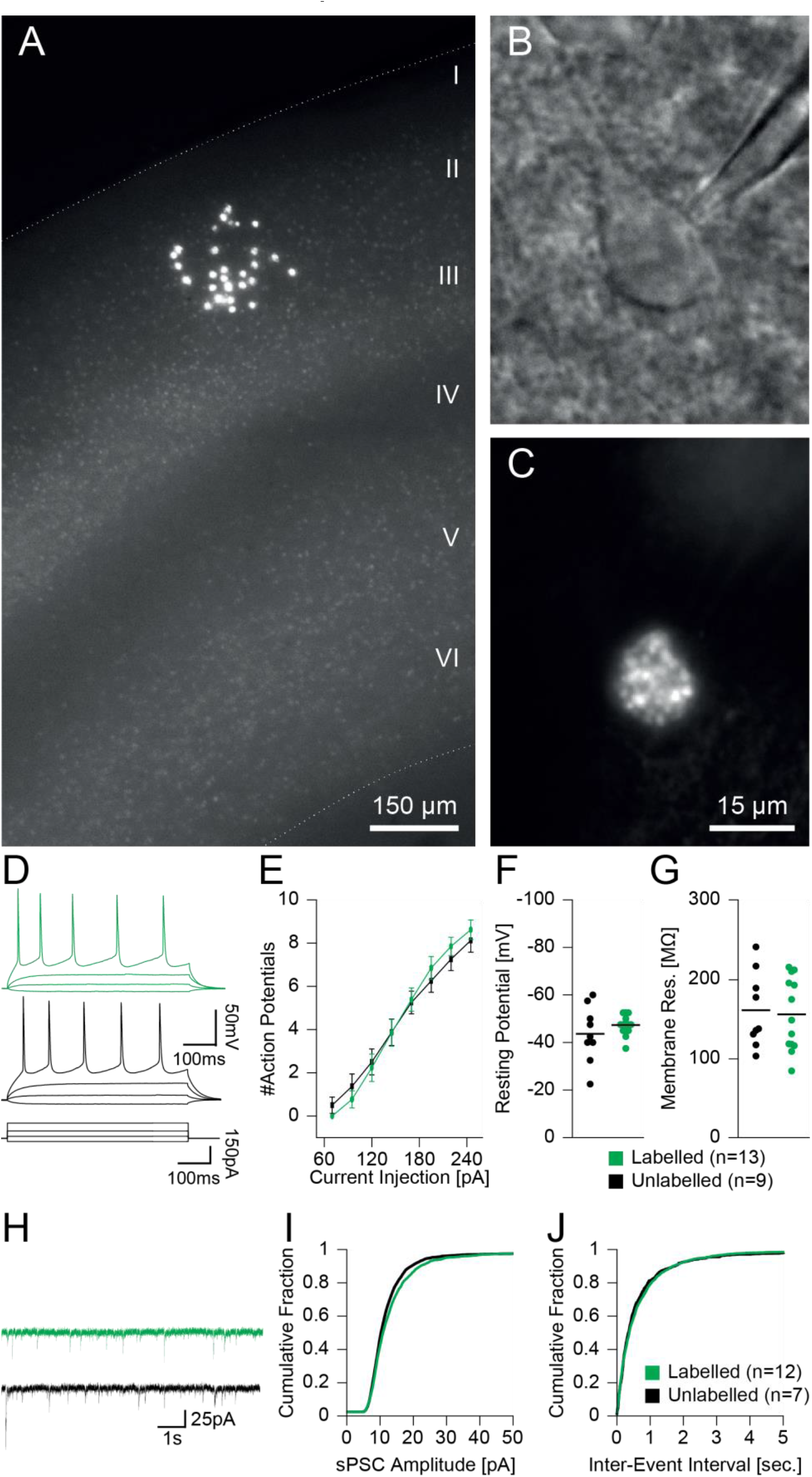
Characterization of electrophysiological properties of photolabeled neurons in transgenic mice with broad forebrain expression of H2B::Pa-GFP. (**A**) Epifluorescence image of an acute coronal brain slice of a H2B::Pa-GFP mouse crossed with a CaMKIIa-Cre mouse with a field of two-photon photolabeled cells in layers II/III. (**B**) Dodt-contrast image of the soma of a pyramidal neuron with a patch-pipette in an acute brain slice. (**C**) Corresponding epifluorescence image of panel (B) showing strong photolabel of the neuronal chromatin. (**D**) Patchclamp whole-cell current-clamp recordings of a photolabeled (green) and non-labeled (black) nearby neuron in response to current injections of various amplitudes. (**E**) Input/output relationship of injected current and elicited action potentials in photolabeled (green) and non-labeled neurons (black). Photolabeled neurons: n=13; 20pA: 0±0.0 AP; 45pA: 0±0.0 AP; 70pA: 0±0.1 AP; 95pA: 0±0.4 AP; 120pA: 1±0.6 AP; 145pA: 4±0.6 AP; 170pA: 5±0.5 AP; 195pA: 7±0.5 AP; 220pA: 8±0.4 AP; 245pA 9±0.4 AP; Non-labeled neurons; n=8; 20pA: 0±0.0 AP; 45pA: 0±0.0 AP; 70pA: 0±0.4 AP; 95pA: 1±0.6 AP; 120pA: 2±0.6 AP; 145pA: 4±0.6 AP; 170pA: 5±0.5 AP; 195pA: 6±0.5 AP; 220pA: 7±0.5 AP; 245pA 8±0.5 AP; all values are median±sem; Two-way ANOVA: significant effect of current injection (d.f.: 9, F=211, p<0.001), no significant effect of photolabeling treatment (d.f.: 1, F=0.009, p=0.93) or interaction (d.f.:9, F=0.91, p=0.53); (**F**) Resting potential in photolabeled (green) and non-labeled neurons (black). Photolabeled neurons: n=13; −79±0.5 mV; Non-labeled neurons: n=9; −77±1.6 mV; p=0.40, Mann-Whitney test. (**G**) Membrane resistance in photolabeled (green) and non-labeled neurons (black). Photolabeled neurons: n=13; 148.2±12.7 MΩ; Non-labeled neurons: n=9; 137.3±15.8 MΩ; p = 0.79, Mann-Whitney test. (**H**) Voltage-clamp registration at −70mV holding potential of spontaneous postsynaptic currents (sPSCs) in photolabeled (green) and non-labeled neurons (black). Photolabeled neurons: n=2021 events, 12 cells; 10.58±0.1 pA; Non-labeled neurons: n=1146 events, 7 cells; 11.23±0.2 pA; p=0.99, Mann-Whitney test). (**I**) Cumulative distributions of sPSC amplitudes in photolabeled (green) and non-labeled neurons (black). (**J**) Cumulative distributions of sPSC inter-event intervals in photolabeled (green) and non-labeled neurons (black).

**Fig. S5:**
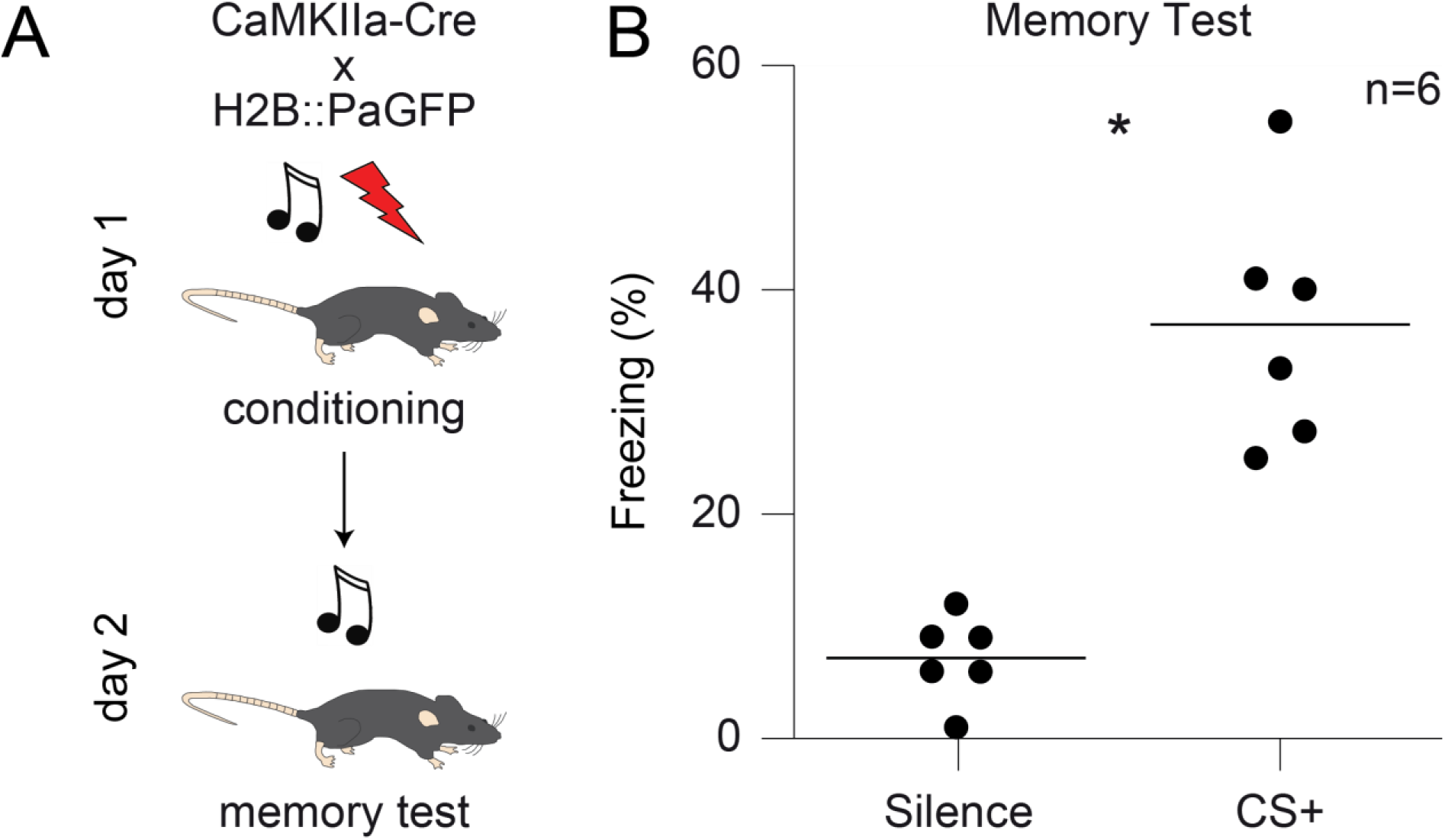
Successful acquisition of fear memories in transgenic mice with broad forebrain expression of H2B::Pa-GFP. **(A)** Offspring of a cross of CaMKIIa-Cre and H2B::Pa-GFP mice underwent auditory cued fear conditioning following a standard protocol involving five sound / foot shock pairings to a complex sound CS+ (see Materials and Methods). **(B)** Quantification of the memory test session 24 hours after auditory cued fear conditioning. Mice showed increased freezing upon CS+ presentation, indicating successful formation of an associative memory (n = 6, Silence: 0.072±0.015, CS+: 0.369±0.045; ANOVA, p<0.001).

**Fig. S6:**
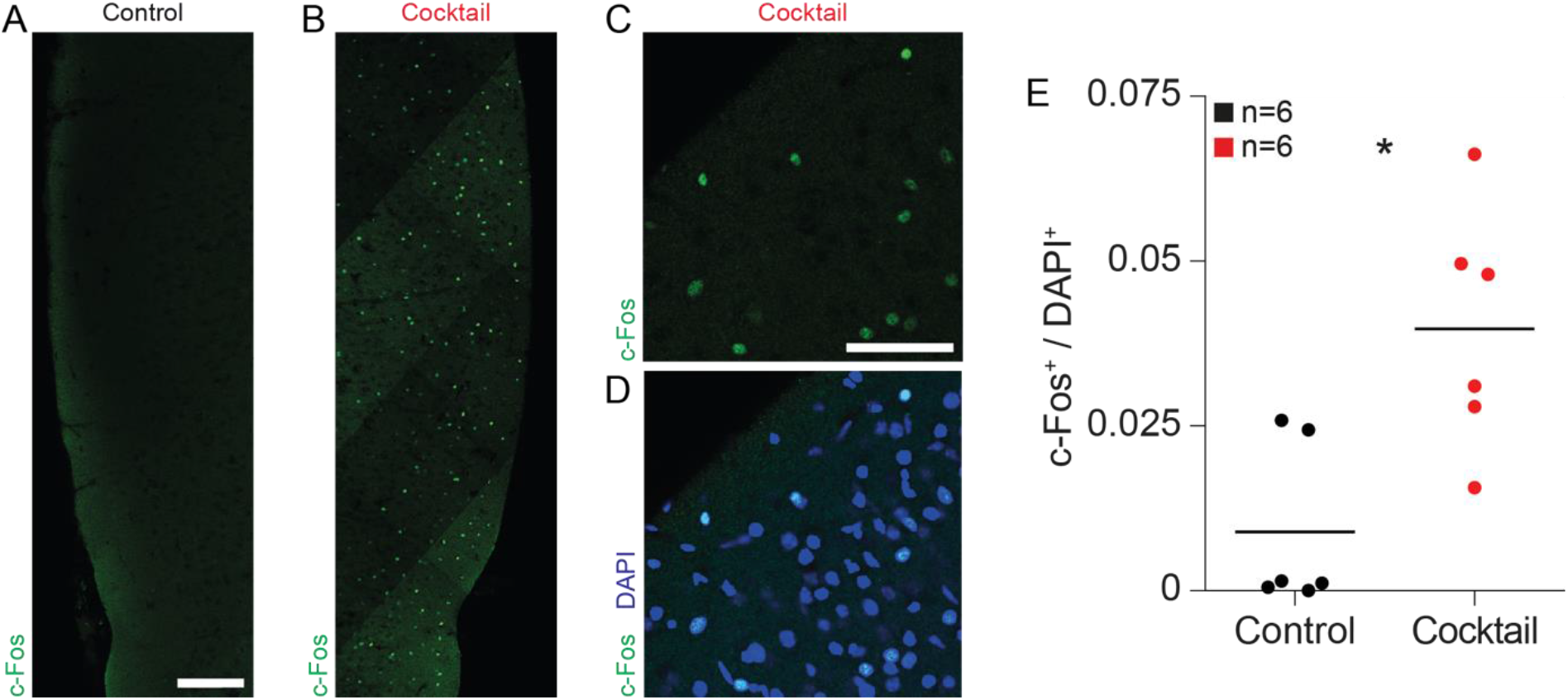
Pharmacological induction of burst activity in acute brain slices leads to increased c-Fos expression. (**A**) Confocal image of a control brain slice fixed 1 hour after sham treatment and subsequent immunofluorescence detection of the IEG product c-Fos. Scale bar: 150 μm. (**B**) Analog image as shown in panel (A), however, brain slice was treated for 1 hour with cocktail. Note increase in the number of c-Fos-positive neurons. (**C**) Higher magnification image of c-Fos-positive cells in a brain slices incubated with cocktail. Scale bar: 50 μm. (**D**) Same image, additionally DAPI staining of nuclei shown in blue. (**E**) Quantification of c-Fos-positive neurons in control brain slices incubated with ACSF (black) and brain slices additionally incubated with cocktail (red). Control: n = 6; 0.13±0.51%; Cocktail: n = 6; 3.95±0.74%; all values are median±sem. Mann-Whitney test: p<0.01.

**Fig. S7:**
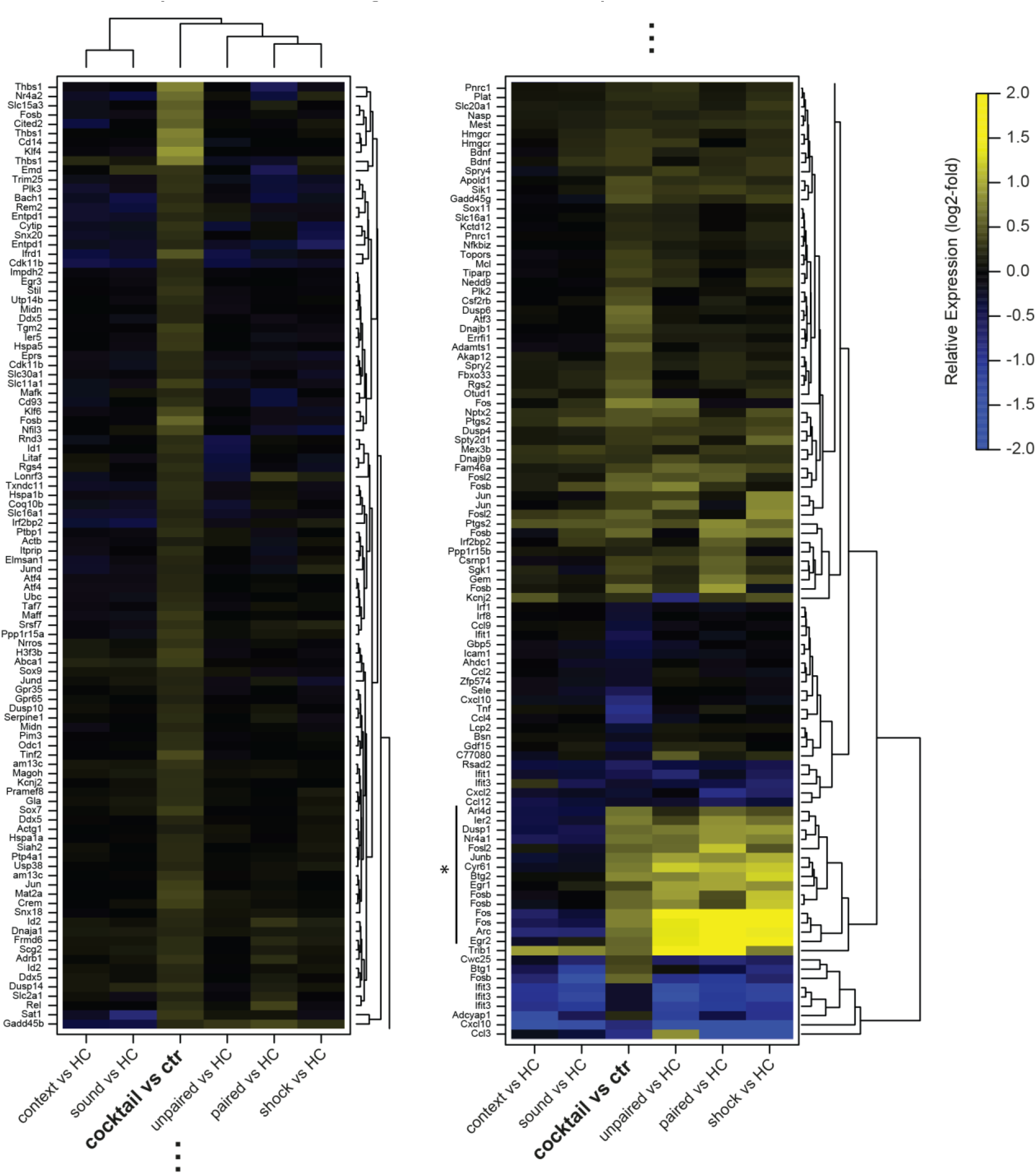
Shared features in gene expression patterns in brain slices after pharmacological induction of neuronal activity and *in vivo* following various behavioral experiences. Heatmap display showing significantly differentially expressed genes analyzed by RNAseq in six acute brain slices treated with the pharmacological cocktail and six control slices (column ‘cocktail vs. control’). In a previous study, we analyzed the effect of various behavioral treatments (paired auditory cued fear conditioning, unpaired auditory cued fear conditioning, shock presentation only, sound presentation only, temporary housing in neutral context, housing in home cage) on gene expression using a microarray-based analysis. Here, we reported, considering the relative expression levels of individual genes, that many known immediate-early genes were induced in those mice undergoing a treatment involving a shock (Peter et al., 2012); EMBL-EBI ArrayExpress, Accession number: E-MTAB-661). For comparison, corresponding relative expression levels from this *in vivo* study are displayed as additional columns. Expression levels of some genes were measured with several probes on the microarray and are therefore mentioned multiple times. Fig. 3, E of the main text of the manuscript shows a subcluster of the heatmap display containing many known immediate-early genes.

**Fig. S8:**
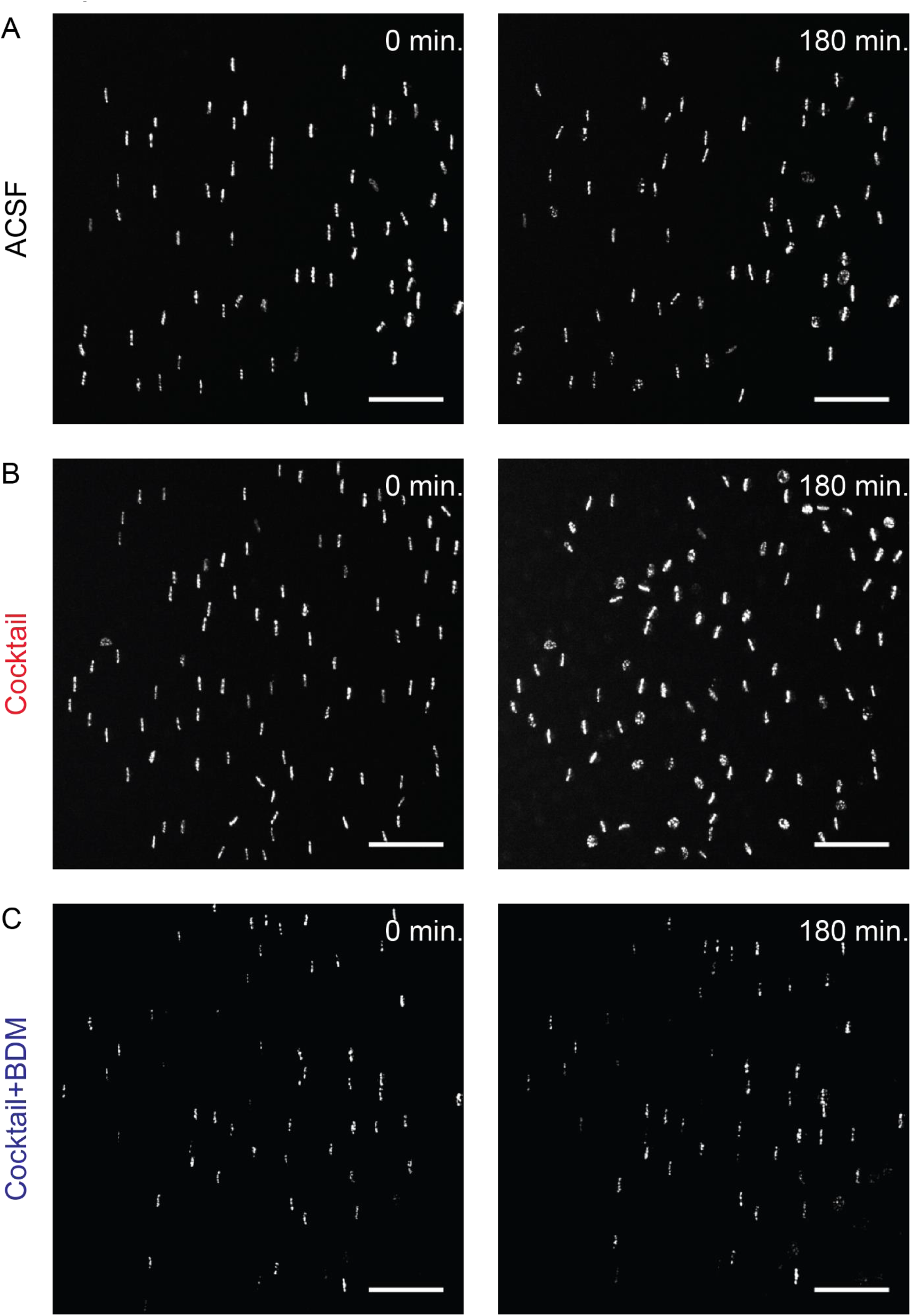
The degree of nucleus-scale reorganization does not correlate with photolabeling efficiency. (**A**) Maximum intensity projection of an experiment with mock manipulation using ACSF. Left: image acquired directly after photolabeling. Right: image acquired 180 minutes after photolabeling. (**B**) Analog to panel (A), but for a slice which was manipulated by exchange of the perfusion solution to the pharmacological cocktail between 60 and 120 minutes after photolabeling. (**C**) Analog to panel (A), but for a slice which was treated with the pharmacological cocktail and BDM.

**Fig. S9:**
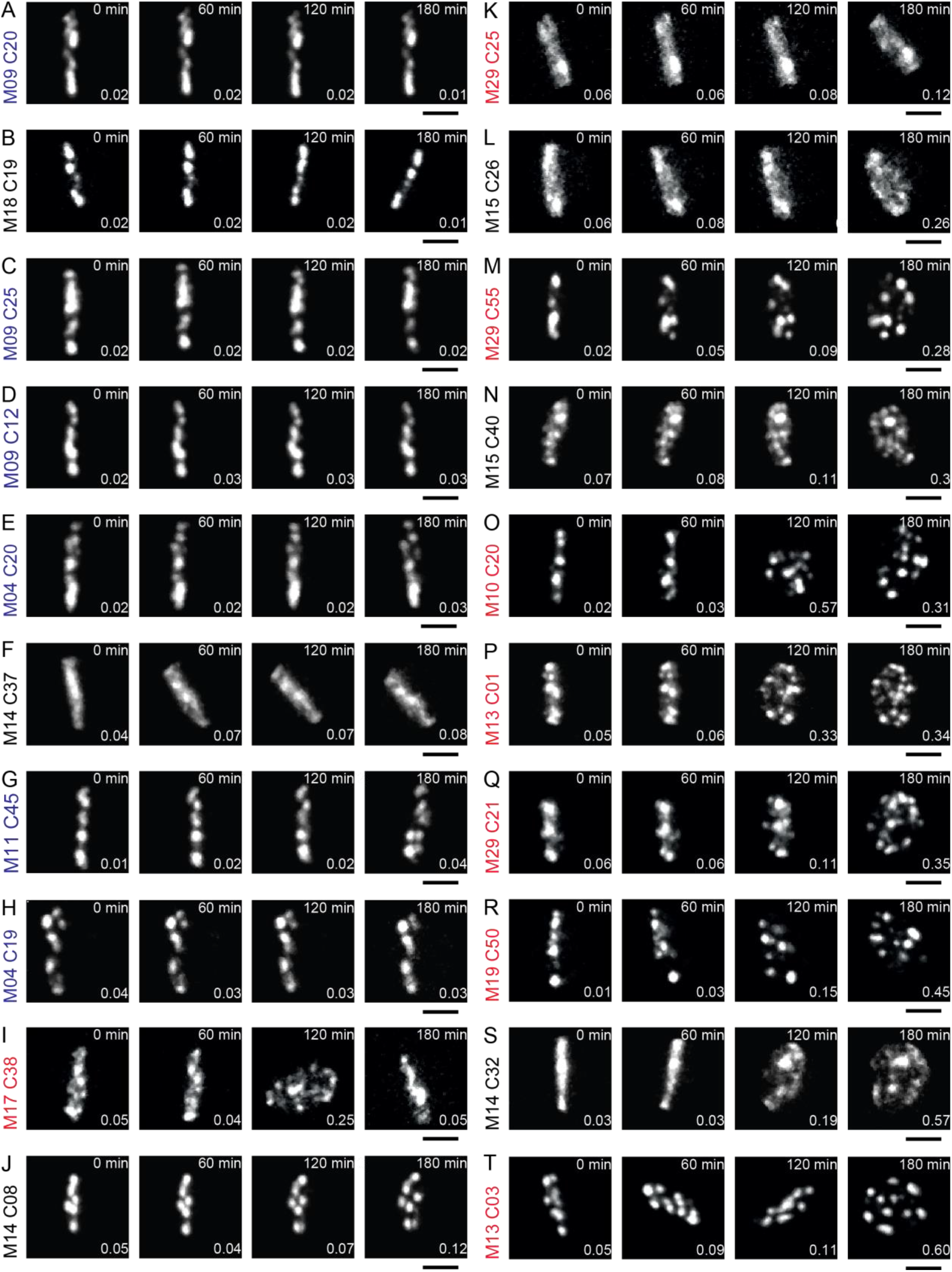
Chromatin dynamics are heterogeneous in brain tissue. Despite clear overall effects between the different treatment conditions (Fig. 3 of the main manuscript), we observed in acute brain slices substantial heterogeneity across individual nuclei within a treatment group. **(A)** to **(T)**: Maximum intensity projections of two-photon image stacks of 20 cells from experiments observing chromatin dynamics in slices from H2B::Pa-GFP crossed with CaMKIIa-Cre mice from all three treatment conditions illustrating heterogeneous chromatin dynamics. Text left of image series depicts identity of cell and slice (Black: Control; Red: Cocktail; Blue: Cockatil+BDM; scale bar = 5 μm).

**Fig. S10:**
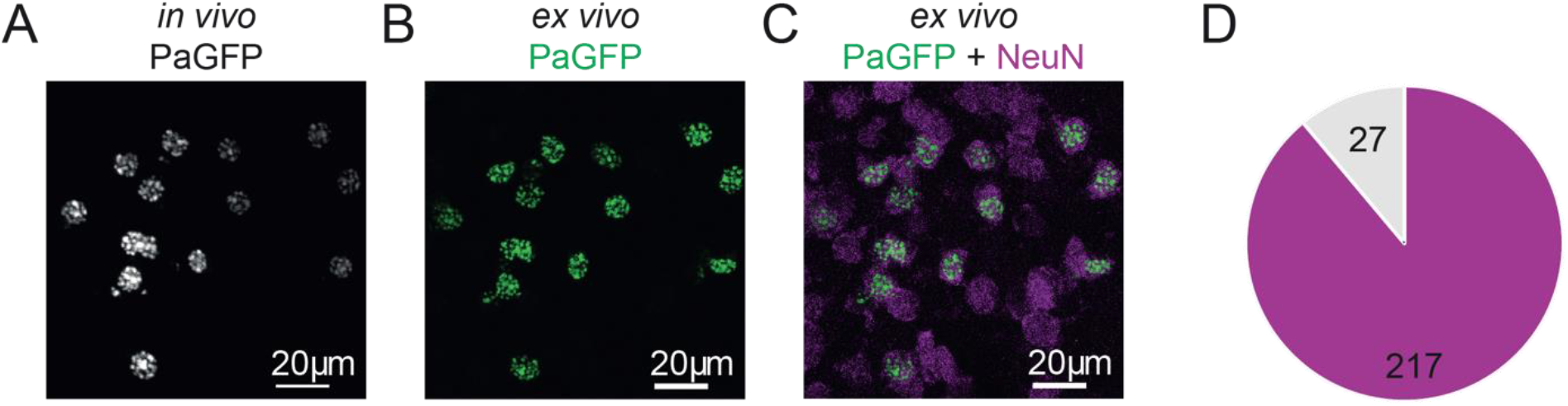
Post-hoc immunofluorescence analysis of photolabeled cells *in*. (**A**) Maximum intensity projection of a *in vivo* two-photon image stack of photolabeled cells in layers II/III of the mouse auditory cortex of a H2B::Pa-GFP mouse with rAAV-mediated Cre-expression under the control of the CMV promoter. We selected for large nuclei for photolabeling that could be identified prior labeling using higher laser intensities at 900nm due to basal fluorescence of Pa-GFP. (**B**) Maximum intensity projection of a confocal image stack of the same cells after fixation and tangential slicing of the brain. Strong labeling of the cells readily enables re-identification. (**C**) Maximum intensity projection of the same stack merged with a second imaging channel for spectrally separated immunofluorescence staining against the neuronal marker protein NeuN in magenta. Most labeled cells show co-staining for NeuN. (**D**) Quantification of the immunofluorescence analysis: Out of a total of 244 *in vivo* photolabeled cells from 7 mice 217 were NeuN positive. The effective transduction of neurons by rAAV5 (Aschauer et al., 2013) and the selection of large nuclei for photolabeling are factors contributing to the strong bias towards labeling neurons.

**Fig. S11:**
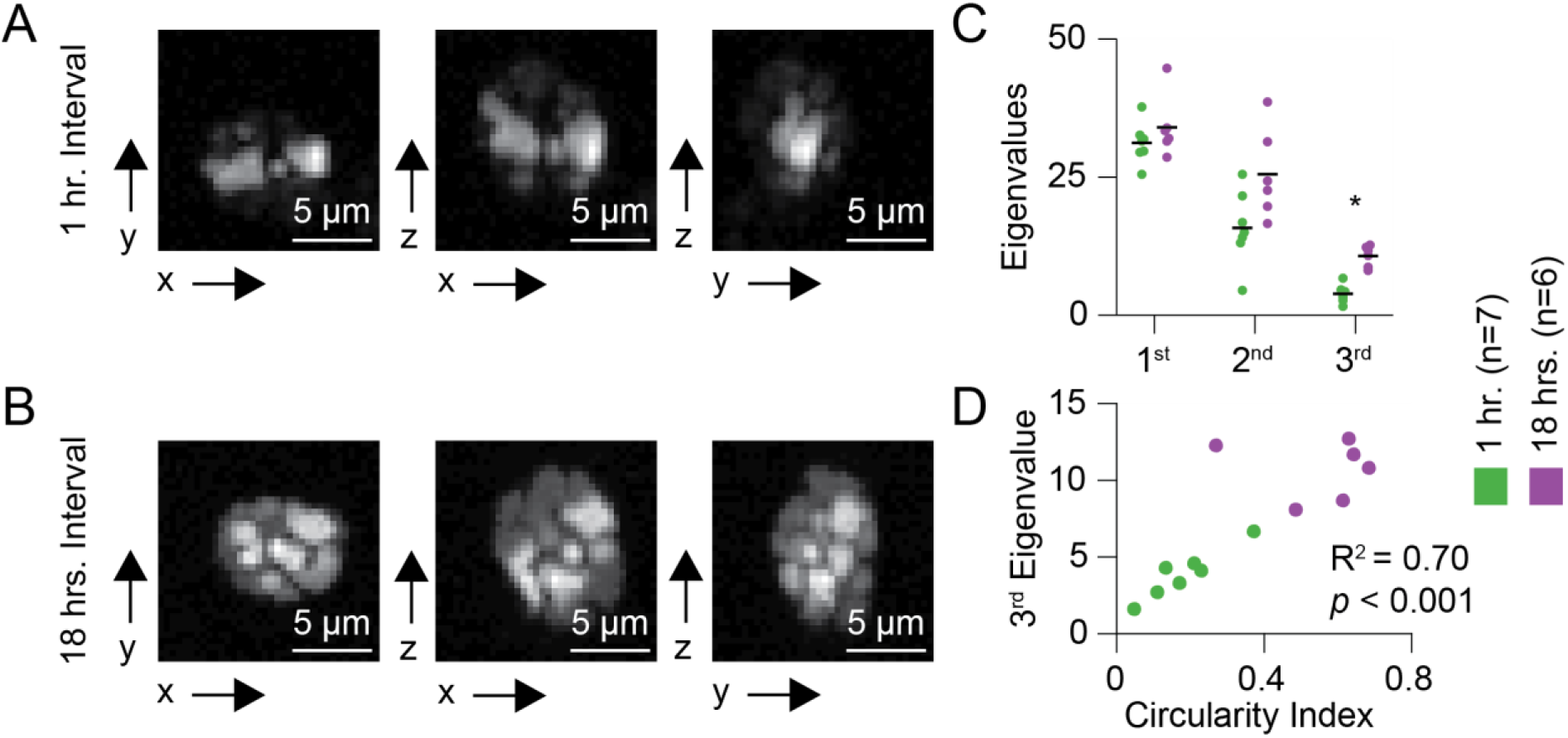
Post-hoc immunofluorescence analysis of photolabeled cells *in vivo* show that chromatin reorganization is reflected by increased CI values. (**A**) Maximum intensity projections of a deconvolved confocal image stack taken from an individual neuron photolabeled 1 hour before fixation. (**B**) Same as panel (A) from an individual neuron photolabeled 18 hours before fixation. Different projections illustrate the spread of the photolabeled subset of chromatin along all three spatial dimensions. (**C**) Three eigenvalues estimated in deconvolved confocal image stacks from neurons fixed 1 hour (red) or 18 hours (blue) after photolabeling. The second and third eigenvalues increase in cells fixed 18 hours after photolabeling indicating spread of the label in all three dimensions. (**D**) Correlation of the CI calculated using the maximum intensity projection along the z-axis and the third eigenvalue estimated in the analysis considering all three dimensions in neurons fixed 1 hour (red) and 18 hours (blue) after photolabeling. In cultured cells, brain slices as well as *in vivo,* we observed rotational movements of the chromatin in many cells. This became particularly obvious in cells in which the photolabeled line was preserved but rotated. This phenotype suggests that nuclear chromatin can move entirely as a rigid body without changing its internal architecture. To test if increases in CI values describing a loss of the line structure are associated with a three-dimensional redistribution of the photolabeled chromatin throughout the nucleus, we performed *in vivo* photolabeling of neurons 18 and 1 hour before fixation, leading to two groups of cells that will, on average, have undergone more or less spontaneous reorganization of chromatin. The photolabeled cells were re-identified in the fixed tissue and imaged using confocal microscopy and subsequent image deconvolution to improve the z resolution of the images. We analyzed the three-dimensional spread of the fluorescence in the image stack using a principal component analysis. Rigid rotational movement would leave the pattern of the photolabel unaltered and, therefore would not affect the eigenvalues describing the spread along the three orthogonal principal components. We observed, however, that the variance along the third and smallest component significantly increased in the group of nuclei photolabeled 18 hours before fixation (1st, 18hrs, n=6, 34.0±2.3, 1hr, n=7, 31.2±1.4, n.s.; 2nd, 18hrs, 25.5±3.3, 1hr, 15.8±2.5, n.s.; 3rd, 18hrs, 10.7±0.8, 1hr, 3.9±0.6, *p*<0.001, unpaired t-test with multiple comparisons). Furthermore, there was a clear correlation between the CI, calculated on the two-dimensional maximum intensity projection along the z-axis, and the third eigenvalue, taking into account the three-dimensional distribution of the photolabel. These results show that, despite clear rotational components in the observed dynamics of chromatin in neuronal nuclei, increases in the CI are correlated with a three-dimensional distribution of the photolabel.

**Fig. S12:**
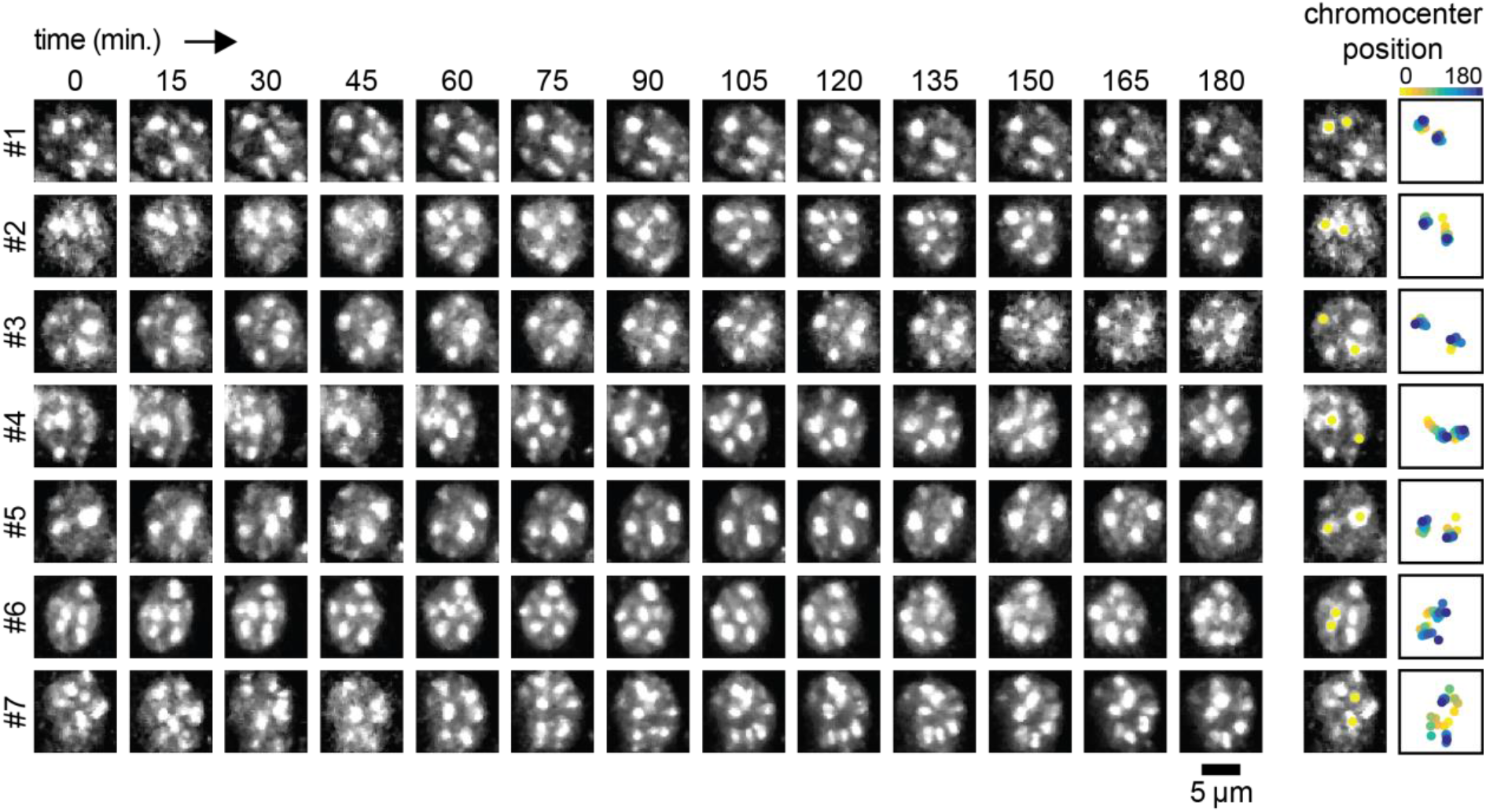
Tracking of chromatin dynamics *in vivo* using bolus-loading of cells with Hoechst. Wild-type C57/Bl6 mice received stereotaxic injections of Hoechst and subsequently a cranial widow was implanted over the auditory cortex. Time-lapse two-photon images (maximum intensity projections) taken from individual nuclei at a 15 min interval during isoflurane anesthesia. Right panels show tracking of the position of two chromocenters per cell over the imaging period with time color coded. Note that, similar to the observations obtained with photolabeling of nucleosomes, different cells show varying levels of chromatin dynamics and phases of relative stability can be interrupted by phases of increased motility.

**Fig. S13:**
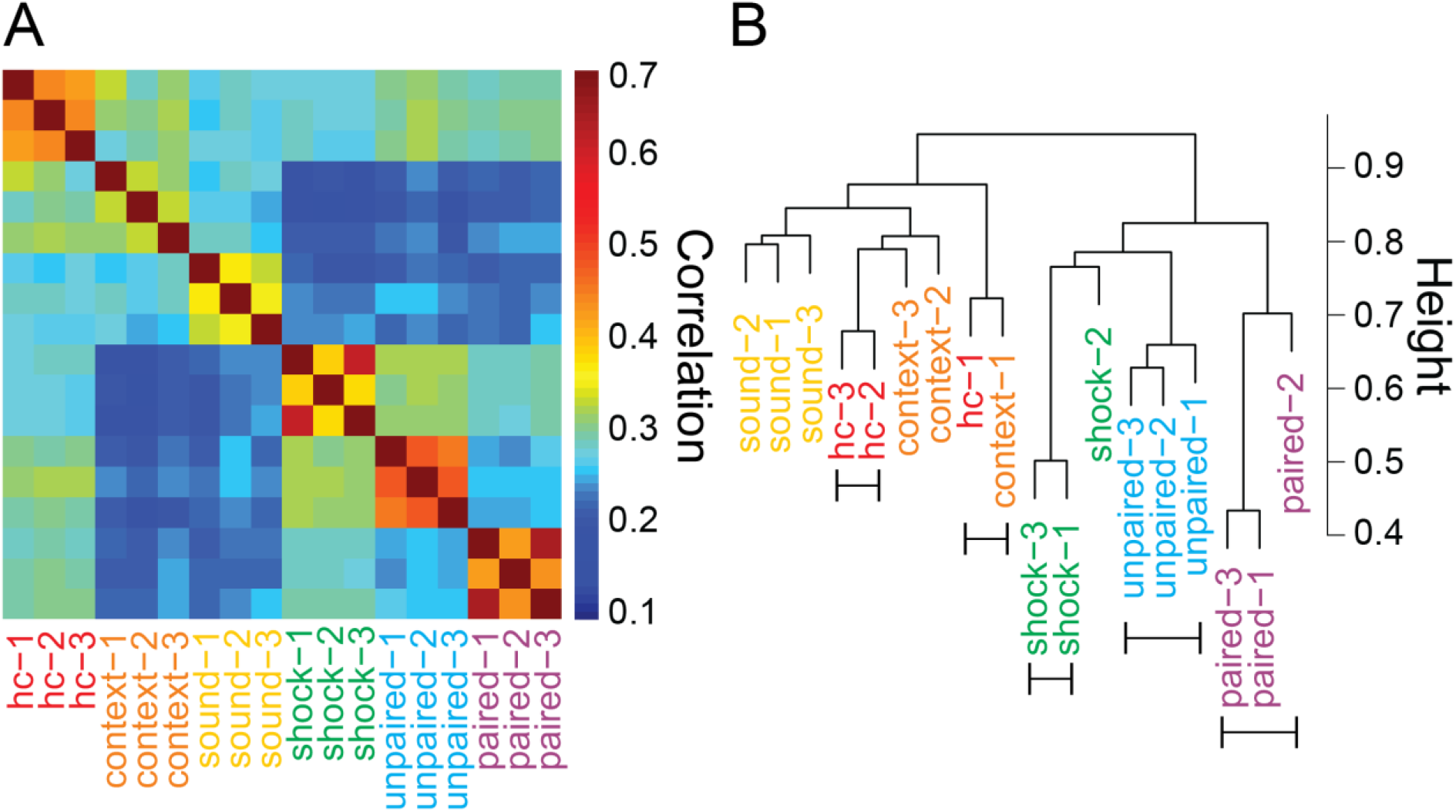
Global gene expression patterns in auditory cortex are specific to behavioral experiences. A re-analysis of the data from the previous microarray study (Peter et al., 2012) taking into account the expression pattern of all genes on the microarray revealed that the biological replicates of mice undergoing either paired or unpaired conditioned could be identified based on the analysis of gene expression patterns. (**A**) Spearman correlation plot of the complete expression profiles, three independent biological replicates per treatment. Paired and unpaired conditioning and to some extend shock presentation only result in a good correlation of their overall expression profile within replicates but is particular distinct from each other as well as from sound presentation and housing in a neutral context. (**B**) Dendrogram of a cluster analysis of the complete expression profiles. The uncertainty in clustering was assessed with the multiscale bootstrap resampling of pvclust (nboot =10000). The bars below indicate clusters with an AU *p*-value > 0.95.

**Fig. S14:**
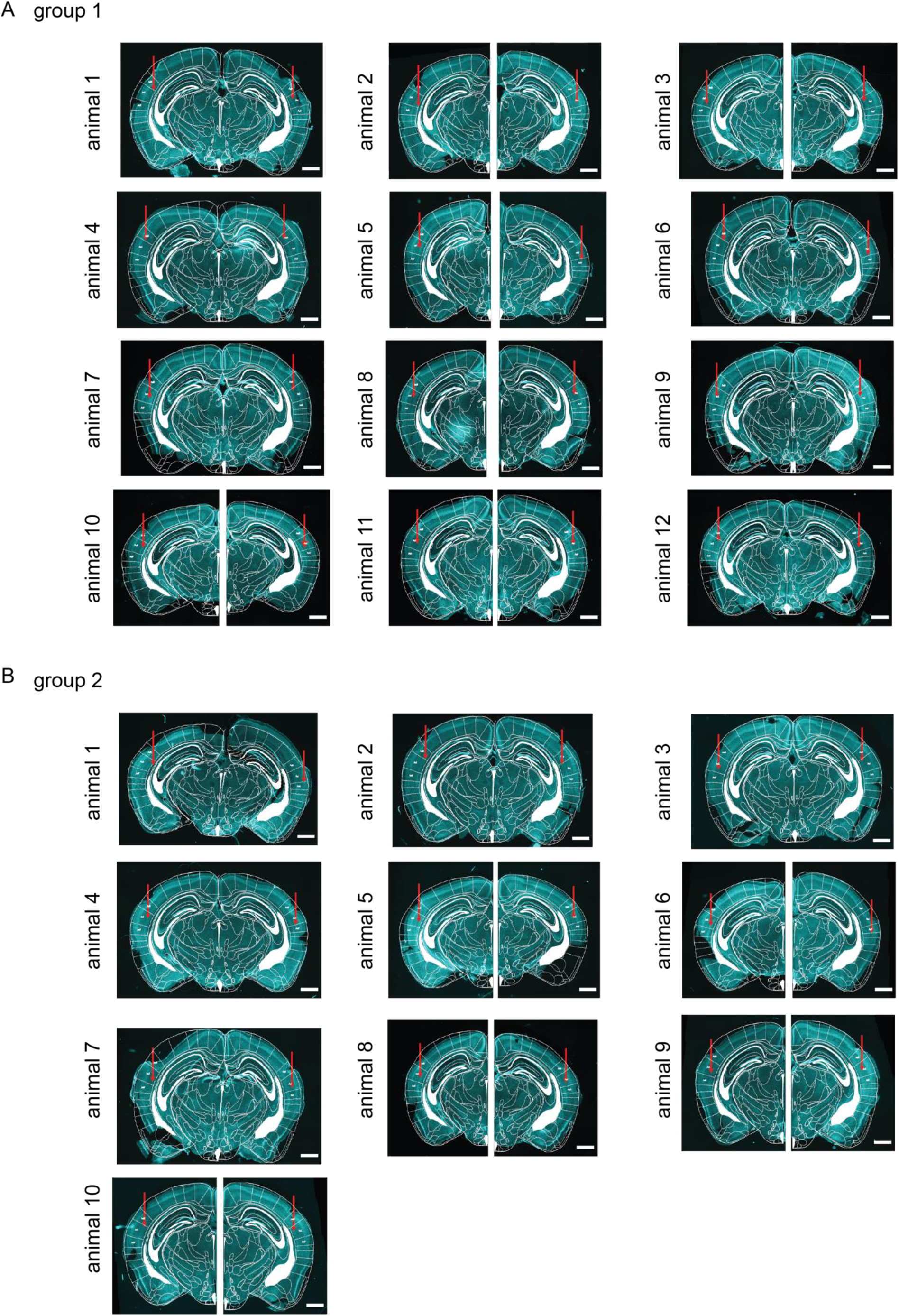
Histological verification of cannulated animals for behavioral impact of BDM during memory acquisition and retrieval. (**A**) Epifluorescence images of DAPI-stained coronal brain sections of mice from group 1 of behavioral experiment to study the impact of bilateral BDM infusion into the auditory cortex. Transparent overlay is from the corresponding section of a mouse brain atlas. Red arrow depicts position of cannula (scale bar = 1 mm). (**B**) Same as panel (A) for group 2 of behavioral experiment.

## Material and methods

### Materials and correspondence

Address correspondence and requests for resources and reagents to corresponding author Simon Rumpel.

### Mice

All animal experiments were performed in accordance with the Austrian laboratory animal law guidelines for animal research and had been approved by the Viennese Magistratsabteilung 58 (Approval: M58/02182/2007/11; M58/02063/2008/8; M58/002220/2011/9). All animal experiments were performed in accordance with institutional guidelines and were approved by the respective Austrian (BGBl nr. 501/1988, idF BGBl I no. 162/2005) and European (Directive 86/609/EEC of 24 November 1986, European Community) authorities and covered by the license GZ2452882016/6.). The mouse line B6.129-Gt(ROSA)26Sortm1(CAG-HIST1H2B/PA-GFP)Rmpl/J will be available from The Jackson Laboratory as Stock No. 035687. It was generated as a further development of the JAX 012071 line for photolableling cells, and shows a substantial increase in the signal/noise of the photolabel, i.e. making the labeled cells much easier detectable and prolonged the longevity of the label. Experimental subjects were both male and female offspring of either homozygous H2B::PaGFP mice or homozygous H2B::PaGFP females crossed with either homozygous B6.129S2-Emx1tm1(cre)Krj/J or B6.Cg-Tg(Camk2a-cre)T29-1Stl/J males.

### Molecular cloning

The sequence of human histone H2B was amplified from a human cDNA library. To obtain the pAAV-CamKIIa-H2B::PaGFP::2A::tdTomato and pAAV-CamKIIa-PaGFP::2A::tdTomato plasmids hChR2(H134R)-EYFP was excised from the pAAV-CaMKIIa-hChR2(H134R)-EYFP plasmid and replaced by H2B::PaGFP::2A::tdTomato or PaGFP::2A::tdTomato using standard molecular cloning techniques. To generate the ROSA26 H2B::PaGFP targeting plasmid the Ai9 plasmid was digested with FseI to remove tdTomato and H2B::PaGFP was inserted into this site to generate Ai9-CAGGS-H2B::PaGFP. This construct contained the 5’ and 3’ homology arms, the CAGGS promoter, a STOP cassette flanked by loxP sites, the H2B::PaGFP transgene, WPRE, polyA and a Neomycin selection cassette.

### Primary dissociated cortical cultures (Fig. 1, 2, S1, S2)

Experiments were performed in primary cortical neurons cultured from newborn (postnatal day 0) mice (C57BL6J) as described previously (Blanquie et al., 2016). After decapitation, brains were transferred to ice-cold Ca^2+^-and Mg^2+^-free HBSS (Gibco, Invitrogen, Carlsbad, CA, USA) supplemented with penicillin and streptomycin (50 units/ml), sodium pyruvate (11 mg/ml), glucose (0.1%), and HEPES (10 mM). Cortical cells were dissociated via trypsin incubation for twenty minutes at 37°C and DNAse digestion at room temperature (RT). After blocking trypsinization by washing steps with HBSS, Minimal Essential Medium (MEM, Gibco) was supplemented with 10% horse serum and 0.6% glucose. Next, cells were mechanically dissociated via repetitive pipetting through fire-polished glass pipettes with declining diameter. Cells were counted after trypan blue staining and seeded with an initial plating density of approximately 1000 cells per mm^2^ on coverslips, on IBIDI dishes (μ-Dish 35mm Grid-500 Glass Bottom, μ-Slide 8 well Grid-500 and μ-Slide 8 well ibiTreat, Ibidi, Germany) or on a multi-electrode array (MEA, Multi channel systems, Reutlingen, Germany). MEAs for electrophysiological recordings were coated with polyethyleneimine (Sigma-Aldrich,Steinheim, Germany; 0.05% in borate-buffered solution). Coverslips and ibidi-dishes were coated with poly-ornithine prior to plating of cells. After 45 minutes, the medium was exchanged for medium consisting of Neurobasal medium (Gibco) supplemented with 2% B27 (Gibco) and 1 mM L-glutamine. Cells were cultivated at 37°C in humidified carbogen (95% air; 5% CO2) for up to 16 days. After two days in vitro (DIV) 5 μM AraC was added to the medium to inhibit glial cell proliferation. Half of the medium was replaced by BrainPhys™ Neuronal Medium (supplemented with SM1 supplement; Stem cell technologies, Vancouver, Canada) every seven days.

### MEA electrophysiological recordings from primary neurons (Fig. S1)

Primary cortical neurons were cultured on MEAs containing 120 planar extracellular titanium nitrite electrodes with four internal references (120MEA100/30iR-Ti-gr, Multi Channel Systems) for 14-16 DIV. For the recordings, signals from 120 recording electrodes were collected with MCRack software in a MEA 2100 system (Multi Channel Systems) at a sampling rate of 50 kHz and high-pass filtered at 200 Hz. Spikes were detected using a threshold-based detector set to a threshold of seven times the standard deviation of the noise level (MC_Rack, Multi Channel Systems). Electrophysiological recordings were performed in culture medium and temperature was maintained at 37°C by a temperature controller (TC02, Multi Channel Systems). Spike datasets from all electrodes were imported into Matlab 7.7 (Mathworks, Natick, MA, USA) for analysis of single units using a custom written routine. Spike sorting was carried out as described previously (Blanquie et al., 2016). Autocorrelation functions were applied to confirm spike sorting. If single units fired at least twice within the recording period, units were counted as active neurons. Average firing frequencies were calculated as arithmetic mean of individual firing frequencies of all identified units and number of active neurons was defined as number of individual single units with at least two action potentials per recording.

### Immunocytochemistry of primary neurons (Fig. 2, S1, S2)

Dissociated cells grown on coverslips were fixed in 2% PFA in PBS for 5 minutes, then transferred to 4% PFA for an additional 15 minutes and then washed with PBS. Unspecific binding of antibodies binding was blocked with 7% normal donkey serum and 0.3% Triton diluted in PBS for two hours at RT. Overnight staining was performed at 4°C with primary antibody diluted in 2% bovine serum albumin with 0.05% azide and 0.1% Triton. Thereafter, cells were washed three times with PBS and incubated with 1 μg/ml DAPI (Sigma-Aldrich), Alexa Fluor647- and DyLight488-coupled secondary antibody (Biomol, Hamburg, Germany) diluted in 2% bovine serum albumin with 0.05% azide in PBS for two hours at RT. Coverslips were washed in PBS and specimens were mounted with Fluoromount (Sigma-Aldrich). The following antibodies were used: Rabbit monoclonal anti c-Fos (Cell Signaling Technology, Inc., SanDiego, CA, USA) 1:800, Rabbit monoclonal anti γH2A.X (phosphor S139) antibody (Abcam, Cambridge, UK) 1:400, guinea pig polyclonal anti c-Fos (Synaptic Systems, Goettingen, Germany) 1:500. Imaging was done on an Olympus IX81 epifluorescence microscope (Olympus Life Sciences, Germany) or on a spinning disc confocal (Microscope (VisiScope 5-Elements, Visitron Systems, Germany) and images were subsequently analyzed with ImageJ or Matlab.

### Live Cell imaging of primary neurons (Fig. 1, 2)

Primary neurons transduced with recombinant AAV carrying either pAAV-hSyn-H2B::mCherry alone or pAAV-hSyn-H2B::mCherry in combination pAAV-hSyn-H2B::PaGFP after one day in culture. Live cell confocal imaging of H2B::mCherry and H2B::PaGFP was performed on DIV 14-16. Images were acquired with a Visiscope 5-Elements spinning disk confocal system (Visitron Systems, Germany), and a 60x water immersion objective (CFI Plan Apo VC 60XWI, 1.2 NA). The system was equipped with a Yokogawa CSU-W1 scan head, a Prime BSI sCMOS camera (2048 x 2048 pixels, 6.5 μm pixel size, Photometrics), a FRAP module that was used for photo-activation of PaGFP, and with an incubator (Bold Line universal stage top incubator, okolab, Italy). Theoretical lateral and axial resolution was 163 nm and 490 nm, respectively, in a field of view of 228 x 228 μm. The laser lines of 488 nm and 561 nm were used for the fluorescence excitation and the fluorescence emission was filtered using filters 525/30 bandpass (Chroma) and 570 longpass (Chroma) for H2B::PaGFP and H2B::mCherry, respectively. The temperature in the incubator was kept at 37°C and the CO2 level at 5% during imaging. Penicillin and streptomycin (5 units/ml) was applied prior to the imaging session to avoid bacterial contamination. Cells were imaged every ten minutes for one hour before the application of pharmacological treatment, and up to two hours after the application. Cells were fixed immediately after the live cell acquisition for immunocytochemical analysis. For experiments with primary cortical neurons co-expressing H2B::mCherry and H2B::PaGFP, photolabeling of a narrow rectangle in the nucleus was performed with a 405 nm laser.

### Calculation of the granularity Index (Fig. 1, 2)

For each image stack of a nucleus, the optical section with the largest signal intensity was self-normalized. A two-dimensional Fourier transform was calculated and a Gaussian function was fitted onto the two cardinal axes of the 2D Fourier transform. The full width half maximum of the average of the two Gaussian fits was termed Granularity Index (GI), which was normalized for each nucleus to its first time point.

### Calculation of the circularity Index (Fig. 2, 3, 4, 5, S6, S7, S8)

To quantify the remodeling of the fluorescence label, we modified a previously established method to measure the geometry of dendritic spines(Moczulska et al., 2013). Briefly, maximum intensity projections of the image stacks of individual nuclei were background subtracted and normalized to maximum intensity values and only intensity values >0.25 were considered for further analysis. A principal component analysis of the 2D pixel map was performed, where each pixel was weighted by its brightness. The circularity index (CI) was defined as the ratio of the smaller to the larger eigenvalue.

### Generation of a H2B::PaGFP knock-in mouse Line (Fig. S3A)

Mitomycin C treated mouse embryonic fibroblasts were plated on 0.1% gelatin coated plates at 37°C, 5% CO2 in DMEM (GIBCO) + 10% FBS, 100 μm Non-essential amino acids, and 2mM L-glutamine. After 24h the medium was exchanged to DMEM + 15% FBS + L-glutamine + 1mM Sodium Pyruvate + 100 μM Non-essential amino acids + 0.1 mM 2-mercaptoethanol + 10^3^ units/ml LIF. A9 129/B6 F1 hybrid embryonic stem cells were grown at 37°C, 5% CO2 and passaged onto mouse embryonic fibroblasts when reaching 90% confluence. The ROSA26 H2B::PaGFP targeting plasmid was linearized using AscI and electroporated into A9 129/B6 F1 hybrid ES cells using standard techniques. Neomycin resistant clones were screened, and positive clones were injected into C57BL6/J blastocysts. Highly chimeric mice were crossed with C57BL/6 mice and successful targeting was confirmed by southern blot. Positive mice were backcrossed to C57BL6/J mice to establish the ROSA26 H2B::PaGFP mouse line.

### Photolabeling in fixed brain slices (Fig. S3B)

To test the specificity of H2B::PaGFP expression and effectiveness of the stop-cassette, H2B::PaGFP mice were crossed with EMX1-Cre mice. Cre-positive and Cre-negative littermates were transcardially perfused with PBS containing 10U/ml Heparin (Sigma-Aldrich) and 4% PFA, the brains removed and post-fixed with 4% PFA overnight at 4°C. Coronal brain slices (70 μm thick) were cut with a vibratome (Leica Biosystems, Germany; VT-1000). Full frame photolabeling and subsequent imaging of PaGFP fluorescence was performed on a LSM780 confocal microscope. The full frame was illuminated using a 405nm laser and subsequent imaging of photolabeled neurons was performed using a 488nm laser.

### Correlation of PaGFP expression and DAPI labeling (Fig. S3C)

H2B::PaGFP mice were injected with a rAAV encoding for Cre recombinase under the control of a CMV promoter. Three weeks later cortical slices were prepared as described above. They were stained with 1 μg/ml DAPI, photolabeled and imaged using an LSM780 confocal microscope.

### Stereotaxic virus injections

Animals were deeply anesthetized with a mixture of ketamine and medetomidine (KM; 2.5 mg ketamine-HCl and 0.02 mg medetomidine-HCl/25 g mouse weight) injected intraperitoneally, and positioned in a stereotaxic frame (Kopf Instruments, Tujunga, CA; Stereotaxic System Kopf 1900). Lidocaine was applied as local anesthetic subcutaneously before exposure of the skull. The virus solution was diluted with PBS to a final titer of 1×1012 GC/ml. Small holes were drilled into the skull and 100 nl were injected at a flow rate of 20 nl/min (World Precision Instruments, Sarasota, FL, USA; Nanoliter 2000 Injector) in five locations along the anterior-posterior axis, resulting in a total injection volume of 500 nl. Stereotactic coordinates were: 4.4, −2,5/-2.75/-3/-3.25/-3.5, 2.5 (in mm, caudal, lateral, and ventral in reference to Bregma). Glass pipettes (World Precision Instruments, Sarasota, FL, USA; Glass Capillaries for Nanoliter 2000) were pulled on a Flaming-Brown Puller (Sutter, Novato, CA, USA) with a long taper and the tip was cut to a diameter of 20-40 μm. After the injection completed, the pipette was left in place for 3 minutes, before being slowly withdrawn and moved to the next coordinate. After completion of the injection protocol, the skin wound was sealed using tissue adhesive (3M Animal Care Products, St. Paul, MN, USA; 3M Vetbond Tissue Adhesive). Mice were monitored daily and intraperitoneal injections of carprofen (0.2 ml of 0.5 mg/ml stock) were applied on the first days after surgery.

### Craniotomy

Male and female H2B::PaGFP mice were used for *in vivo* imaging experiments and a small imaging window was implanted over the auditory cortex three weeks after virus injection. Briefly, mice were deeply anesthetized with a single dose of ketamine/medetomidine. After removal of an about 1 cm^2^ patch of skin over the parietal bones on the right lateral part of the head, a small part of the musculus temporalis was removed to expose the temporal bone. Using a dentist’s drill, the bones were smoothened, and part of the zygomatic process was removed and covered with a thin layer of Vetbond glue (3M). Next, a thin layer of dental cement (Lang Dental) was applied, except for the area over the temporal bone. The temporal bone was removed with a dental drill to expose a ~2 mm × 3 mm part of the brain containing the auditory cortex. The craniotomy was subsequently sealed with a drop of liquid agarose, a small round cover glass, and additional dental cement. In addition, a small titanium head post was implanted for fixation of the head during imaging.

### In vivo two-photon imaging (Fig. 4, 5, S3D)

In vivo imaging and photolabeling was performed using an Ultima *in vivo* multiphoton microscopy system (Prairie Technologies, WI, USA) with a 20x objective lens (Olympus, Tokyo, Japan; XLUMPlan FI, n.a.=0.95) and a Chameleon Ultra Ti:Sapphire multiphoton laser (Coherent, CA, USA). Mice were anesthetized with isofluorane (Abbot Animal Health, IL, USA; IsoFlo) and H2B::PaGFP positive cells were identified by their weak basal fluorescence. For photolabeling of the whole nucleus, a ROI was placed over the soma. Photolabeling was performed at 750nm and imaging of photolabeled cells at 950nm. Photolabeling was done on a single focal plane and images were collected as z-stacks. To estimate the long-term stability of the photolabel *in vivo,* a ROI was placed over the soma of the neuron and the neuron was photolabeled. Individual labeled neurons were revisited and imaged at different time points at the indicated intervals. For analysis, images were background subtracted and the fluorescence was normalized to the fluorescence level measured directly after photolabeling.

### Acute brain slice preparation and electrophysiology (Fig. 3 A – D, S4)

4-week-old mice were deeply anesthetized using isoflurane (Abbot Animal Health, IL, USA; IsoFlo), decapitated and their brains quickly chilled in ice-cold oxygenated (95% O2/ 5% CO2) dissection buffer (110 mM choline chloride, 25 mM NaHCO3, 1.25 mM NaH2PO4, 2.5 mM KCl, 0.5 mM CaCl2, 7 mM MgCl2, 11.6 mM ascorbic acid, 3.1 mM pyruvic acid, and 25 mM D-glucose). Coronal brain slices (350μm) containing auditory cortex were cut in ice-cold dissection buffer using a Vibratome (Leica Biosystems, Germany; VT1000S) and immediately incubated for 15 min in dissection buffer at 32°C. This was followed by a phase in which slices rested in oxygenated (95% O2/5% CO2) artificial cerebrospinal fluid (aCSF; 118 mM NaCl, 2.5 mM KCl, 26.5 mM NaHCO3, 1 mM NaH2PO4, 1 mM MgCl2, 2 mM CaCl2, and 20 mM D-glucose) for 45min at room temperature (RT). Individual brain slices were placed on the stage of an upright, infrared-differential interference contrast microscope (Olympus BX50WI) mounted on a X-Y table (Luigs & Neumann, Ratingen, Germany) and visualized with a 40x water immersion objective by an infrared sensitive digital camera (Hamamatsu, ORCA-03). Slices were fully submerged and continuously perfused at a rate of 1 −2 ml per min with oxygenated aCSF. Pyramidal neurons from superficial layers of auditory cortex were identified by their characteristic shape or, in the case of photolabeled neurons, by GFP fluorescence. Patch pipettes were pulled on a Flaming/Brown micropipette puller (P-97; Sutter, Novato, CA, USA) from borosilicate glass (1.5 mm outer and 0.86 mm inner diameter; Sutter, Novato, CA, USA) to final resistances ranging from 3 to 5 MΩ. Internal solution for voltage and current clamp experiments contained (in mM): 135 K-Gluconate, 5 KCl, 10 HEPES, 2 MgCl2, 0.2 EGTA, 1 Na2ATP, 0.4 NaGTP, 10 Na2Phosphocreatine. Membrane currents were recorded with a Multicamp 700B amplifier (Molecular Devices, Sunnyvale, CA, USA). Electrophysiological signals were low pass filtered at 3 kHz, sampled at 10 kHz (Axon Instruments, Molecular Devices, Sunnyvale, CA, USA; Digidata 1440A) and stored on a PC for offline analysis with pClamp 10 software (Molecular Devices, Sunnyvale, CA, USA). Whole-cell patch clamp recordings from pyramidal neuron pairs in the upper layers of auditory cortex were recorded simultaneously. After 10 minutes of baseline recording, aCSF in the recording bath chamber was replaced by ‘cocktail aCSF’ containing aCSF as described above and in addition 50μM (-)-Bicuculline methiodide, 100μM DL-Norepinephrin hydrochloride, 50μM Carbamoylcholine chloride, 100μM Dopamine hydrochloride, 40μM Ascorbic Acid and 100μM Serotonin hydrochloride (all from Sigma-Aldrich, MO, USA). For experiments investigating incubation with pharmacological cocktail, coronal brain slices containing auditory cortex were prepared as described above. After the initial 15 minutes of recovery in Choline Chloride, the two hemispheres were separated by a cut along the midline under a dissection microscope and incubated in aCSF for 45 minutes. One hemisphere of each brain slice was then transferred to fresh aCSF for 1 hour; the other hemisphere was incubated in cocktail aCSF for 1h. Both, control aCSF and cocktail aCSF hemispheres, were finally exposed to another resting phase in bubbled aCSF for 1 hour. Solutions for each step described above were oxygenated with 95% O2/5% CO2 at 32°C.

### c-Fos immunohistochemistry in coronal brain slices (Fig. S6)

c-Fos/DAPI staining was performed on coronal brain slices incubated in normal aCSF or cocktail aCSF as described above. After fixation in 4% PFA at 4°C for 2 hours, free floating slices were permeabilized with 0.2% Triton X-100 and blocked with 2% normal goat serum and 2% BSA for 2 hours at room temperature. Slides were then incubated with a primary antibody against c-Fos (1:1000, rabbit; Abcam, Cambridge, UK; ab7963) for 24 hours at 4°C, followed by washing and incubation with an Alexa-488 conjugated secondary antibody (1:1000, goat, Invitrogen, CA, USA; A11008) against rabbit at room temperature. Labeled probes were cover slipped with DAPI-Fluoromount-GTM (Biomedica, Vienna, Austria). Images were acquired using the Zeiss LSM 700 point scanning confocal microscope. For detection and quantification of c-Fos signal, a semi-automated, machine learning based approach was implemented with the Pannoramic Viewer image processing software (3D Histech, Budapest, Hungary; Developer XD, Definiens, Munich, Germany). The DAPI signal was used as nuclear reference for the immunodetection of (nuclear) c-Fos. Nuclei were segmented based on the DAPI channel using a LoG-Filter and watershed algorithms. On a set of training images, nuclei objects were manually classified as positive and negative and used as samples for training a machine-learning algorithm. Mean intensity, standard deviation and local contrast parameters were extracted from these as input for a decision tree-based classifier. This classifier was then applied to all nuclei objects on a larger set of images to identify positive cells. A similar approach was used to segment and classify cytoplasmic stain around the nuclei. Training was done separately for each channel. The output was given as number of positive cells in each channel, either c-Fos or DAPI, co-staining of cells, total cell number and mean intensity per cell for further analysis. To quantify differences in cumulative fluorescence probability of c-Fos between control and cocktail samples, average c-Fos fluorescence values from each DAPI-stained nucleus were normalized to the individual background of each image using R (R foundation, Vienna, Austria). Fluorescence data from all mice in each group were pooled to generate an empirical cumulative distribution function (ECDF) for each group.

### Imaging chromatin dynamics in coronal brain slices (Fig. 3, F – K, S8)

An individual slice was transferred to a slice recording chamber in an Ultima multiphoton microscopy system (Prairie Technologies, WI, USA), gently immobilized by a silver grid with attached nylon mesh, and fully submerged and continuously perfused with oxygenated aCSF at room temperature. Imaging was performed using a 20x objective lens (XLUMPlan FI, 0.95 N.A., Olympus) and a Ti:Sapphire multiphoton laser (Coherent, CA, USA). Neurons of layer II/III were identified by baseline PaGFP fluorescence at 950 nm, and were labeled by applying 750 nm laser light on a rectangular ROI over the nucleus at a single focal plane. After labeling of a population of neurons, z stacks were acquired at 0.5 μm steps. Imaging parameters were: 1.2 us pixel dwell time, 2048×2048 pixel resolution at 0.1344 microns per pixel. Photolabeled populations of neurons were revisited and imaged every hour for six hours. After one hour of aCSF perfusion, experimental slices were perfused with aCSF containing 50μM (-)-Bicuculline methiodide, 100μM DL-Norepinephrin hydrochloride, 50μM Carbamoylcholine chloride, 100μM Dopamine hydrochloride, 40μM Ascorbic Acid and 100μM Serotonin hydrochloride (all Sigma-Aldrich, MO, USA) for one hour and returned to normal aCSF for the rest of the experiment. Slices in the Cocktail + BDM group were treated with 20 mM 2,3-butanedione monoxime (Sigma) in addition to the described pharmacological cocktail. Control slices were perfused with normal aCSF during the whole course of the experiment including a ‘mock switch’ at the same timing as the experimental groups. Individual labeled neurons were manually identified in z-stacks of different time points and isolated by cropping of ROIs of 100×100 pixels. Maximum intensity z-projections were used for calculation of the CI using the same custom written Matlab (Mathworks, Natick, MA, USA) script as for the *in vivo* data.

### RNA preparation, sequencing, and expression analysis (Fig. 3E, S7)

Brain slice hemispheres that underwent incubation protocol in cocktail or control aCSF, as described above, were placed under a dissection microscope and auditory cortices dissected while submerged in bubbled aCSF. Total RNA from dissected auditory cortices was isolated by Trizol (800μl / auditory cortex; Life technologies, Grand Island, NY, USA). Total RNA was quantified and quality assessed with a Bioanalyzer (Agilent Technologies, CA, USA) RNA 6000 Nano kit. One microgram of total RNA was used for poly-A selection with a Dynabeads mRNA purification kit (Invitrogen, CA, USA) followed by cDNA generation (NEB 1st strand and NEB 2nd strand ultra directional Kits; New England BioLabs, Frankfurt am Main, Germany). Libraries were prepared using a NEBNext Ultra DNA kit (New England BioLabs, Frankfurt am Main, Germany) and quantified with a Library Quantification kit for Illumina platforms (Kapa Biosystems, Inc., Wilmington, MA, USA). Sequencing was performed on a HiSeq 2500 system (Illumina, San Diego, CA, USA) following the manufacturer’s guidelines in single read modus and at a read length of 50bp. All samples were multiplexed in one lane returning 15 to 27 million reads with an average Q score above 30 per sample. The strand specific reads were screened for ribosomal

RNA by aligning with BWA (v0.6.1) against known rRNA sequences (RefSeq). The rRNA subtracted reads were aligned with TopHat (v1.4.1) against the Mus musculus genome (mm10) and a maxiumum of 6 missmatches. Maximum multihits was set to 1 and InDels as well as Microexon-search was enabled. Additionally, a gene model was provided as GTF (UCSC RefSeq mm10). rRNA loci are masked on the genome for downstream analysis. Aligned reads are subjected to FPKM estimation with Cufflinks (v1.3.0). In this step bias detection and correction was performed. Furthermore, only those fragments compatible with UCSC RefSeq annotation (mm10) of genes with at least one protein coding transcript were allowed and counted towards the number of mapped hits used in the FPKM denominator. Furthermore, the aligned reads were counted with HTSeq (0.6.1p1) and the genes were subjected to differential expression analysis with DESeq2 (v1.6.3; Bioconductor). The effect of samples originating from the same mouse was blocked.

### Auditory cued fear conditioning and memory testing (Fig. 6, S5)

Behavioral experiments were performed in an isolation cubicle (Coulbourn Instruments, Whitehall, PA, USA) which was equipped with white LEDs as house light, a microphone and a CCD KB-R3138 camera with infrared LEDs (LG Electronics Austria, Vienna, Austria) which was connected to a Cronos frame grabber (Matrox, Dorval, Quebec, Canada). The conditioning chamber (25 × 25 × 42 cm, Coulbourn Instruments) was combined either with a stainless steel shock floor or a grid floor. A custom-made cartridge (round or quadrangular) was inserted to form different local environmental contexts. Foot shocks were delivered via an external shocker (Precision Animal shocker, Coulbourn Instruments). Sounds were played from a L-22 soundcard with a maximal sampling frequency of 192 kHz (Lynx Studio Technology, Costa Mesa, CA,USA) and delivered via an amplifier (Model SLA-1, Applied Research and Technology, TEAC Europe GmbH, TASCAM Division, Wiesbaden, Germany), a modified equalizer (Model #351, Applied Research and Technology, TEAC Europe GmbH, TASCAM Division, Wiesbaden, Germany) and a custom-made speaker for free field delivery of sounds. Sound levels for all stimuli used were normalized to a mean power of 78 dB sound pressure level (SPL). Peak sound levels ranged from 83 to 89 dB SPL. For auditory cued fear conditioning five sound-foot shock pairings (0.75 mA, 1 second, immediately following the sound) were delivered with a randomized inter-stimulus time interval ranging from 50 to 75 seconds. For unpaired conditioning, five foot shocks and five sounds were presented in a randomized order separated by at least one minute. Memory testing was performed 24 hours after conditioning in a different context and consisted of five sound presentations. Videos of the mice were recorded during the memory test and analyzed using a custom Matlab (Mathworks, Natick, MA, USA) script calculating the relative time spent freezing. Freezing behavior was automatically scored based on movement rate (frame-to-frame difference, significant motion pixels detected on the movies). Baseline freezing was assessed during silence between 30 and 60 seconds of each protocol run.

### Intrinsic imaging (Fig. 5B)

Mice were lightly anesthetized with isofluorane (Abbot Animal Health, IL, USA; IsoFlo) and the area under the cranial window was imaged with a 780nm light emitting diode and a CCD camera (Vosskühler, Osnabrück, Germany). Sound stimuli consisted of white noise, pure tones and the complex sound used for auditory cued fear conditioning. Sounds were presented 30 times and the change in light reflectance was computed and averaged using custom Matlab (Mathworks, Natick, MA, USA) software.

### Imaging chromatin dynamics *in vivo* (Fig. 4, 5F)

For in vivo chromatin imaging, large nuclei in layer II/III were identified for labeling using PaGFP baseline fluorescence. Photolabeling was performed sound responsive areas, as determined via intrinsic imaging. For photolabeling, a 5x zoom factor was used (x,y-pixel size 0.215 x 0.215 microns) and a small ROI in the pattern of a thin stripe was placed over the nucleus. Then a single focal plane was illuminated with 750nm leading to a thin fluorescent rod in the nucleus. Photolabeled cells were imaged as z-stacks (1um step size) in duplicates.To analyze the three-dimensional spread of fluorescence and to assess the three-dimensional pattern in labeled cells, lines were photolabeled and brains fixated after 1h or 18h. When performing photolabeling using two-photon excitation, the minimal extent of the label is determined by the point-spread function of the focus. The point spread function is affected by the specific imaging conditions *in vivo* and is typically in the submicron range in the xy-dimension, whereas it extends to about 2 μm along the z-dimension. Thus, the extent of the photolabeled lines along the z-axis is likely larger than their lateral extent. As in many cells a clear rotational component became apparent, we wondered if the transformation of the line pattern to a more homogeneous labeling of the nucleus can be accounted for by a rigid 3D rotation of the whole chromatin along the x or y axis. Labeled nuclei were re-identified in fixed brain slices and were imaged at high magnification using an LSM780 confocal microscope. Images were deconvolved using Huygens software (Scientific Volume Imaging (SVI), Hilversum, Netherlands). After deconvolution, image stacks of individual nuclei were re-sampled to obtain an isotropic resolution of 0.368 μm for all three dimensions. Again, pixel intensities were background subtracted and normalized to maximum intensity values. A 2D principal component analysis was performed on a maximum intensity projection as described above. To assess the three-dimensional pattern a principal component analysis of the 3D pixel map was performed considering intensity values >0.1 and the eigenvalues calculated describing the spread of fluorescence along the three principal axes.

### Tracking of chromatin dynamics *in vivo* using bolus-loading of cells with Hoechst (Fig. S12)

A craniotomy was performed on 8-week-old mice and 200nl of a 10 μg/ml solution of Hoechst 33342 (ThermoFisher) was injected in multiple spots in 200 – 300 μm depth. Image stacks of stained cells were acquired in 15 minutes intervals for three hours in 200 – 300 μm depth at 750 nm wavelength.

### Chromatin photolabeling and behavior (Fig. 5, D – H)

Mice were habituated for 3 days prior to the experiment. This included handling by the experimenter, placing into the behavior chambers, anaesthesia with isoflurane (Abbot Animal Health, IL, USA; IsoFlo) and head fixation in the two-photon microscope. The behavior and imaging paradigm lasted for five days (anesthesia, context, unpaired, paired, and memory test). Anesthesia (day one): Mice were anesthetized and placed into the two-photon microscope. Nuclei were labeled with a line pattern and z-stacks of the population were acquired (Pre). The mice remained under the microscope and the same neurons were imaged again two hours later (Post). Context (day two): Mice were anesthetized and placed into the two-photon microscope. A different group of neurons from the first day were labeled with a stripe and z-stacks of the population were acquired (Pre). Afterwards, the mice were removed from the microscope, put back into their cage and allowed to wake up. Two hours after the first imaging session, mice were anesthetized, placed back under the microscope and stacks of the same ensemble of neurons were recorded (Post). Unpaired (day three): Mice were anesthetized and placed into the two-photon microscope. Another ensemble of nuclei was labeled with a stripe and imaged (Pre). Afterwards the mice were put back into their cage and allowed to wake up. Then the mice were subjected to one unpaired conditioning session and put back in their cage. Two hours after the first imaging session, mice were anesthetized and put back into the microscope and stacks of the same ensemble were recorded (Post). Paired (day four): Mice were anesthetized and placed under the two-photon microscope. Another ensemble of nuclei was labeled with a stripe and z-stacks of the population were recorded (Pre). Afterwards, the mice were put back into their cage and allowed to wake up. Then the mice were subjected to one paired conditioning session and put back in their cage. Two hours after the first imaging session, mice were anesthetized and put back under the two-photon microscope and stacks of the same neurons were recorded (Post). 24 hours after the paired session, the acquisition of an associative memory was assessed in a memory test session (day five).

### Guide cannula implantation and infusion combined with auditory cued fear conditioning and memory testing (Fig. 6)

Male 2-4 month old C57BL/6J mice (Charles River) were stereotaxically implanted with 26GA guide cannulas bilaterally (PlasticsOne) 1mm above infusion target area under isoflurane anesthesia. Cannulas were mounted on the skull with dental cement (Super-Bond C&B, Sun Medical) and animals let recover for 1 week. Animals received analgesics (Carprofen, Rimadyl) via drinking water postsurgery. Bilateral infusion of 250nl aCSF (40nl/min each) or 250nl 2,3-butanedione monoxime (BDM, 20 mM) via a 33GA infusion cannula (PlasticsOne) by an infusion pump (Harvard apparatus) was done 20 minutes prior behavioral experiments. All behavioral experiments were performed in an isolation cubicle which was equipped with white LEDs as house light and a CCD camera (Basler acA1300-30gc) with infrared LEDs. The conditioning chamber (17 × 17 × 42 cm) was combined with either a grid floor (context A) or a cylindrical inset and a stainless-steel shock grid (context B). Mice were habituated in a context A and context B on day 1. On day 2 mice underwent auditory cued fear conditioning 1 in context B after infusion of aCSF (Group 1) or BDM (Group 2), where each of 5 sound presentations (2s, 70dB) was immediately followed by a foot-shock (0.75 mA, 1 second; Precision Animal shocker, Coulbourn Instruments) with randomized inter-stimulus time intervals ranging from 50 to 75 seconds. On day 3 memory was tested in context A (recall 1) by 2 blocks of the 20s sound (10x 2s) after infusion of aCSF to both groups. On day 4 mice were re-conditioned (conditioning 2) in context B after aCSF infusion to both groups, followed by a memory test (recall 2) after infusion with inverted group treatments, compared to conditioning 1, on day 5. Freezing was tracked in Observer v11 (Noldus) by an experimenter blind to the treatment. Freezing values are given as % time of the duration of all freezing bouts during the respective experimental periods (baseline and CS periods). Baseline freezing was assessed during silence between 30 and 60 seconds of each protocol run.

### Post hoc immunohistochemistry of *in vivo* photolabeled neurons (Fig. S10)

After the photolabeling session, mice were transcardially perfused as described with a two-hour postfix protocol in 4% PFA. 70 μm thick slices were cut on a vibratome (VT-1000, Leica) parallel to the imaging plane. Free floating sections were incubated for 2 hours at room temperature in PBS containing 10% normal goat serum (NGS) and 1% Triton-X 100. They were washed with PBS and incubated overnight at 4°C in PBS with 5% NGS, 0.1% Triton-X 100 and an anti NeuN antibody (raised in guinea pig). On the next day, sections were washed three times with PBS for ten minutes at room temperature and incubated for two hours at room temperature in PBS containing 5% NGS, 0.1% Triton-X 100 and an anti-guinea pig secondary antibody coupled to Alexa Fluor 647. Next, sections were washed three times with PBS for 10 minutes at room temperature and coversliped. Photolabeled cells could be readily identified in the slice by eye based on their strong fluorescence signal. Confocal stacks were obtained on a LSM 780 confocal laser scanning microscope (Zeiss) and co-localisation was analyzed using ImageJ.

### Statistical analysis

The statistical analyses were performed using Graphpad Prism 6 (Graphpad Software, La Jolla, CA, USA) and Matlab (Mathworks, Natick, MA, USA). We used an appropriate statistical test for the given statistical analysis. For the data to analyze three dimensional distribution of the chromatin label, we used an unpaired t-test with multiple comparisons. For the comparison of the immunohistochemical analysis of c-Fos expression in brain slices, we used a Mann-Whitney test. For all comparisons of the circularity index between groups and time points, we used a two-way ANOVA with a post-hoc Sidak’s multiple comparison test. All values are given as mean±sem unless otherwise noted. Statistical parameters including the exact value of n, precision measures and statistical significance are reported in the text, the figures, and the figure legends. The significance threshold was placed at α=0.05 (n.s., p>0.05; *, p<0.05).

### Histological Verification of Cannula Position (Fig. S14)

Mice were transcardially perfused with 4% PFA in PBS. The brains were extracted, and fixed with 4% PFA over night at 4 °C. The next day, brains were coronally sliced and stained with DAPI, as described above. Epifluorescence images of the section with the largest extent of cannula damage were acquired with an AF7000 Widefield Fluorescence Microscope (Leica, Germany) and an N PLAN 5x/0.12 DRY objective to estimate the position of the cannula post-hoc. The final position of the cannula in the tissue was estimated from the surface of the brain and the length of the cannula and animals with a mis-hit were excluded (group 1: 12/16 included, group 2: 10/16 included).

### Data Availability

RNASeq data of brain slices incubated with a pharmacological cocktail

GEO accession number: GSE94826

Reviewer link: https://www.ncbi.nlm.nih.gov/geo/query/acc.cgi?token=qnmnysciprwvfen&acc=GSE94826

## References

Abraham, W.C., Mason, S.E., Demmer, J., Williams, J.M., Richardson, C.L., Tate, W.P., Lawlor, P.A., and Dragunow, M. (1993). Correlations between immediate early gene induction and the persistence of long-term potentiation. Neuroscience 56, 717–727.

Aschauer, D.F., Kreuz, S., and Rumpel, S. (2013). Analysis of Transduction Efficiency, Tropism and Axonal Transport of AAV Serotypes 1,2, 5, 6, 8 and 9 in the Mouse Brain. Plos One 8, e76310.

Banani, S.F., Lee, H.O., Hyman, A.A., and Rosen, M.K. (2017). Biomolecular condensates: organizers of cellular biochemistry. Nature Reviews Molecular Cell Biology 18, 285–298.

Barr, M.L., and Bertram, E.G. (1949). A Morphological Distinction between Neurones of the Male and Female, and the Behaviour of the Nucleolar Satellite during Accelerated Nucleoprotein Synthesis. Nature 163, 676–677.

Beagan, J.A., Pastuzyn, E.D., Fernandez, L.R., Guo, M.H., Feng, K., Titus, K.R., Chandrashekar, H., Shepherd, J.D., and Phillips-Cremins, J.E. (2020). Three-dimensional genome restructuring across timescales of activity-induced neuronal gene expression. Nature Neuroscience 23, 707–717.

Belyaeva, A., Venkatachalapathy, S., Nagarajan, M., Shivashankar, G.V., and Uhler, C. (2017). Network analysis identifies chromosome intermingling regions as regulatory hotspots for transcription. Proc Natl Acad Sci U S A 114, 13714–13719.

Bertaina, V., and Destrade, C. (1995). Differential time courses of c-fos mRNA expression in hippocampal subfields following acquisition and recall testing in mice. Cognitive Brain Research 2, 269–275.

Bickmore, W.A., and van Steensel, B. (2013). Genome Architecture: Domain Organization of Interphase Chromosomes. Cell 152, 1270–1284.

Blanquie, O., Liebmann, L., Hübner, C.A., Luhmann, H.J., and Sinning, A. (2016). NKCC1-Mediated GABAergic Signaling Promotes Postnatal Cell Death in Neocortical Cajal-Retzius Cells. Cerebral Cortex 27, 1644–1659.

Bohn, M., Diesinger, P., Kaufmann, R., Weiland, Y., Muller, P., Gunkel, M., von Ketteler, A., Lemmer, P., Hausmann, M., Heermann, D.W., et al. (2010). Localization microscopy reveals expressiondependent parameters of chromatin nanostructure. Biophys J 99, 1358–1367.

Boija, A., Klein, I.A., Sabari, B.R., Dall’Agnese, A., Coffey, E.L., Zamudio, A.V., Li, C.H., Shrinivas, K., Manteiga, J.C., Hannett, N.M., et al. (2018). Transcription Factors Activate Genes through the PhaseSeparation Capacity of Their Activation Domains. Cell 175, 1842–+.

Borden, J., and Manuelidis, L. (1988). Movement of the X-Chromosome in Epilepsy. Science 242, 1687–1691.

Buchwalter, A., Kaneshiro, J.M., and Hetzer, M.W. (2019). Coaching from the sidelines: the nuclear periphery in genome regulation. Nature Reviews Genetics 20, 39–50.

Cavalli, G., and Misteli, T. (2013). Functional implications of genome topology. Nature Structural & Molecular Biology 20, 290–299.

Chambeyron, S., and Bickmore, W.A. (2004). Chromatin decondensation and nuclear reorganization of the HoxB locus upon induction of transcription. Genes & Development 18, 1119–1130.

Chang, L., Godinez, W.J., Kim, I.H., Tektonidis, M., de Lanerolle, P., Eils, R., Rohr, K., and Knipe, D.M. (2011). Herpesviral replication compartments move and coalesce at nuclear speckles to enhance export of viral late mRNA. Proceedings of the National Academy of Sciences of the United States of America 108, E136–E144.

Cho, W.K., Spille, J.H., Hecht, M., Lee, C., Li, C., Grube, V., and Cisse, II (2018). Mediator and RNA polymerase II clusters associate in transcription-dependent condensates. Science 361, 412–415.

Chuang, C.-H., and Belmont, A.S. (2007). Moving chromatin within the interphase nucleus-controlled transitions? Seminars in Cell & Developmental Biology 18, 698–706.

Chuang, C.H., Carpenter, A.E., Fuchsova, B., Johnson, T., de Lanerolle, P., and Belmont, A.S. (2006). Long-range directional movement of an interphase chromosome site. Current Biology 16, 825–831.

Chubb, J.R., Boyle, S., Perry, P., and Bickmore, W.A. (2002). Chromatin motion is constrained by association with nuclear compartments in human cells. Current Biology 12, 439–445.

Cremer, T., and Cremer, C. (2001). Chromosome territories, nuclear architecture and gene regulation in mammalian cells. Nature Reviews Genetics 2, 292–301.

Csink, A.K., and Henikoff, S. (1998). Large scale chromosomal movements during interphase progression in Drosophila. Journal of Cell Biology 143, 13–22.

De Boni, U., and Mintz, A. (1986). Curvilinear, three-dimensional motion of chromatin domains and nucleoli in neuronal interphase nuclei. Science 234, 863–866.

Dekker, J., and Mirny, L. (2016). The 3D Genome as Moderator of Chromosomal Communication. Cell 164, 1110–1121.

Dekker, J., and Misteli, T. (2015). Long-Range Chromatin Interactions. Cold Spring Harbor Perspectives in Biology 7, a019356.

Dion, V., and Gasser, S.M. (2013). Chromatin Movement in the Maintenance of Genome Stability. Cell 152, 1355–1364.

Dion, V., Kalck, V., Horigome, C., Towbin, B.D., and Gasser, S.M. (2012). Increased mobility of double-strand breaks requires Mec1, Rad9 and the homologous recombination machinery. Nature Cell Biology 14, 502–U221.

Fischer, A. (2014). Epigenetic memory: the Lamarckian brain. EMBO J 33, 945–967.

Fomproix, N., and Percipalle, P. (2004). An actin-myosin complex on actively transcribing genes. Experimental Cell Research 294, 140–148.

Gasser, S.M. (2002). Nuclear architecture – Visualizing chromatin dynamics in interphase nuclei. Science 296, 1412–1416.

Gibson, B.A., Doolittle, L.K., Schneider, M.W.G., Jensen, L.E., Gamarra, N., Henry, L., Gerlich, D.W., Redding, S., and Rosen, M.K. (2019). Organization of Chromatin by Intrinsic and Regulated Phase Separation. Cell 179, 470–484.

Gu, B., Swigut, T., Spencley, A., Bauer, M.R., Chung, M., Meyer, T., and Wysocka, J. (2018). Transcription-coupled changes in nuclear mobility of mammalian cis-regulatory elements. Science 359, 1050–1055.

Gutnick, M.J., Connors, B.W., and Prince, D.A. (1982). Mechanisms of neocortical epileptogenesis in vitro. Journal of Neurophysiology 48, 1321–1335.

Guzowski, J.F., McNaughton, B.L., Barnes, C.A., and Worley, P.F. (1999). Environment-specific expression of the immediate-early gene Arc in hippocampal neuronal ensembles. Nat Neurosci 2, 1120–1124.

Halder, R., Hennion, M., Vidal, R.O., Shomroni, O., Rahman, R.U., Rajput, A., Centeno, T.P., van Bebber, F., Capece, V., Vizcaino, J.C.G., et al. (2016). DNA methylation changes in plasticity genes accompany the formation and maintenance of memory. Nature Neuroscience 19, 102–+.

Heale, V., and Harley, C. (1990). MK-801 and AP5 impair acquisition, but not retention, of the Morris milk maze. Pharmacology Biochemistry and Behavior 36, 145–149.

Hnisz, D., Shrinivas, K., Young, R.A., Chakraborty, A.K., and Sharp, P.A. (2017). A Phase Separation Model for Transcriptional Control. Cell 169, 13–23.

Hubner, M.R., and Spector, D.L. (2010). Chromatin Dynamics. Annual Review of Biophysics, Vol 39 39, 471–489.

Kanda, T., Sullivan, K.F., and Wahl, G.M. (1998). Histone-GFP fusion protein enables sensitive analysis of chromosome dynamics in living mammalian cells. Curr Biol 8, 377–385.

Korzus, E., Rosenfeld, M.G., and Mayford, M. (2004). CBP histone acetyltransferase activity is a critical component of memory consolidation. Neuron 42, 961–972.

Kosak, S.T., Skok, J.A., Medina, K.L., Riblet, R., Le Beau, M.M., Fisher, A.G., and Singh, H. (2002). Subnuclear compartmentalization of immunoglobulin loci during lymphocyte development. Science 296, 158–162.

Lanctot, C., Cheutin, T., Cremer, M., Cavalli, G., and Cremer, T. (2007). Dynamic genome architecture in the nuclear space: regulation of gene expression in three dimensions. Nat Rev Genet 8, 104–115.

Larson, A.G., Elnatan, D., Keenen, M.M., Trnka, M.J., Johnston, J.B., Burlingame, A.L., Agard, D.A., Redding, S., and Narlikar, G.J. (2017). Liquid droplet formation by HP1alpha suggests a role for phase separation in heterochromatin. Nature 547, 236–240.

Levenson, J.M., O’Riordan, K.J., Brown, K.D., Trinh, M.A., Molfese, D.L., and Sweatt, J.D. (2004). Regulation of histone acetylation during memory formation in the hippocampus. Journal of Biological Chemistry 279, 40545–40559.

Loewenstein, Y., Kuras, A., and Rumpel, S. (2011). Multiplicative Dynamics Underlie the Emergence of the Log-Normal Distribution of Spine Sizes in the Neocortex In Vivo. Journal of Neuroscience 31, 9481–9488.

Madabhushi, R., Gao, F., Pfenning, A.R., Pan, L., Yamakawa, S., Seo, J., Rueda, R., Phan, T.X., Yamakawa, H., Pao, P.C., et al. (2015). Activity-Induced DNA Breaks Govern the Expression of Neuronal Early-Response Genes. Cell 161, 1592–1605.

Marco, A., Meharena, H.S., Dileep, V., Raju, R.M., Davila-Velderrain, J., Zhang, A.L., Adaikkan, C., Young, J.Z., Gao, F., Kellis, M., et al. (2020). Mapping the epigenomic and transcriptomic interplay during memory formation and recall in the hippocampal engram ensemble. Nature Neuroscience.

Marshall, W.F., Straight, A., Marko, J.F., Swedlow, J., Dernburg, A., Belmont, A., Murray, A.W., Agard, D.A., and Sedat, J.W. (1997). Interphase chromosomes undergo constrained diffusional motion in living cells. Current Biology 7, 930–939.

Martou, G., and De Boni, U. (2000). Nuclear topology of murine, cerebellar Purkinje neurons: Changes as a function of development. Experimental Cell Research 256, 131–139.

Masui, O., Bonnet, I., Le Baccon, P., Brito, I., Pollex, T., Murphy, N., Hupe, P., Barillot, E., Belmont, A.S., and Heard, E. (2011). Live-Cell Chromosome Dynamics and Outcome of X Chromosome Pairing Events during ES Cell Differentiation. Cell 145, 447–458.

Maze, I., Wenderski, W., Noh, K.M., Bagot, R.C., Tzavaras, N., Purushothaman, I., Elsasser, S.J., Guo, Y., Ionete, C., Hurd, Y.L., et al. (2015). Critical Role of Histone Turnover in Neuronal Transcription and Plasticity. Neuron 87, 77–94.

Mehta, I.S., Amira, M., Harvey, A.J., and Bridger, J.M. (2010). Rapid chromosome territory relocation by nuclear motor activity in response to serum removal in primary human fibroblasts. Genome Biol 11, R5.

Miller, C.A., Gavin, C.F., White, J.A., Parrish, R.R., Honasoge, A., Yancey, C.R., Rivera, I.M., Rubio, M.D., Rumbaugh, G., and Sweatt, J.D. (2010). Cortical DNA methylation maintains remote memory. Nat Neurosci 13, 664–666.

Moczulska, K.E., Tinter-Thiede, J., Peter, M., Ushakova, L., Wernle, T., Bathellier, B., and Rumpel, S. (2013). Dynamics of dendritic spines in the mouse auditory cortex during memory formation and memory recall. Proceedings of the National Academy of Sciences of the United States of America 110, 18315–18320.

Muratani, M., Gerlich, D., Janicki, S.M., Gebhard, M., Eils, R., and Spector, D.L. (2002). Metabolic-energy-dependent movement of PML bodies within the mammalian cell nucleus. Nature Cell Biology 4, 106–110.

Neumann, F.R., Dion, V., Gehlen, L.R., Tsai-Pflugfelder, M., Schmid, R., Taddei, A., and Gasser, S.M. (2012). Targeted INO80 enhances subnuclear chromatin movement and ectopic homologous recombination. Genes Dev 26, 369–383.

Patterson, G.H., and Lippincott-Schwartz, J. (2002). A photoactivatable GFP for selective photolabeling of proteins and cells. Science 297, 1873–1877.

Peleg, S., Sananbenesi, F., Zovoilis, A., Burkhardt, S., Bahari-Javan, S., Agis-Balboa, R.C., Cota, P., Wittnam, J.L., Gogol-Doering, A., Opitz, L., et al. (2010). Altered Histone Acetylation Is Associated with Age-Dependent Memory Impairment in Mice. Science 328, 753–756.

Peter, M., Bathellier, B., Fontinha, B., Pliota, P., Haubensak, W., and Rumpel, S. (2013). Transgenic Mouse Models Enabling Photolabeling of Individual Neurons In Vivo. Plos One 8, e62132.

Peter, M., Scheuch, H., Burkard, T.R., Tinter, J., Wernle, T., and Rumpel, S. (2012). Induction of immediate early genes in the mouse auditory cortex after auditory cued fear conditioning to complex sounds. Genes Brain and Behavior 11, 314–324.

Ricci, M.A., Manzo, C., Garcia-Parajo, M.F., Lakadamyali, M., and Cosma, M.P. (2015). Chromatin fibers are formed by heterogeneous groups of nucleosomes in vivo. Cell 160, 1145–1158.

Riccio, A. (2010). Dynamic epigenetic regulation in neurons: enzymes, stimuli and signaling pathways. Nature Neuroscience 13, 1330–1337.

Yuen, R.K.C, Merico, D., Bookman, M., Howe, J.L., Thiruvahindrapuram, B., Patel, R.V., Whitney, J., Deflaux, N., Bingham, J., Wang, Z., et al. (2017). Whole genome sequencing resource identifies 18 new candidate genes for autism spectrum disorder. Nat Neurosci 20, 602–611.

Sabari, B.R., Dall’Agnese, A., Boija, A., Klein, I.A., Coffey, E.L., Shrinivas, K., Abraham, B.J., Hannett, N.M., Zamudio, A.V., Manteiga, J.C., et al. (2018). Coactivator condensation at super-enhancers links phase separation and gene control. Science 361, eaar3958–3913.

Sanulli, S., Trnka, M.J., Dharmarajan, V., Tibble, R.W., Pascal, B.D., Burlingame, A.L., Griffin, P.R., Gross, J.D., and Narlikar, G.J. (2019). HP1 reshapes nucleosome core to promote phase separation of heterochromatin. Nature 575, 390–394.

Schoenfelder, S., Sexton, T., Chakalova, L., Cope, N.F., Horton, A., Andrews, S., Kurukuti, S., Mitchell, J.A., Umlauf, D., Dimitrova, D.S., et al. (2010). Preferential associations between coregulated genes reveal a transcriptional interactome in erythroid cells. Nature Genetics 42, 53–U71.

Shrinivas, K., Sabari, B.R., Coffey, E.L., Klein, I.A., Boija, A., Zamudio, A.V., Schuijers, J., Hannett, N.M., Sharp, P.A., Young, R.A., et al. (2019). Enhancer Features that Drive Formation of Transcriptional Condensates. Molecular Cell 75, 549–+.

Strickfaden, H., Zunhammer, A., van Koningsbruggen, S., Kohler, D., and Cremer, T. (2010). 4D chromatin dynamics in cycling cells Theodor Boveri’s hypotheses revisited. Nucleus 1, 284–297.

Strom, A.R., Emelyanov, A.V., Mir, M., Fyodorov, D.V., Darzacq, X., and Karpen, G.H. (2017). Phase separation drives heterochromatin domain formation. Nature 547, 241–245.

Suberbielle, E., Sanchez, P.E., Kravitz, A.V., Wang, X., Ho, K., Eilertson, K., Devidze, N., Kreitzer, A.C., and Mucke, L. (2013). Physiologic brain activity causes DNA double-strand breaks in neurons, with exacerbation by amyloid-beta. Nature Neuroscience 16, 613–+.

Szabo, Q., Bantignies, F., and Cavalli, G. (2019). Principles of genome folding into topologically associating domains. Science Advances 5, eaaw1668.

Thakar, R., and Csink, A.K. (2005). Changing chromatin dynamics and nuclear organization during differentiation in Drosophila larval tissue. Journal of Cell Science 118, 951–960.

Therizols, P., Illingworth, R.S., Courilleau, C., Boyle, S., Wood, A.J., and Bickmore, W.A. (2014). Chromatin decondensation is sufficient to alter nuclear organization in embryonic stem cells. Science 346, 1238–1242.

van Steensel, B., and Belmont, A.S. (2017). Lamina-Associated Domains: Links with Chromosome Architecture, Heterochromatin, and Gene Repression. Cell 169, 780–791.

Vazquez, J., Belmont, A.S., and Sedat, J.W. (2001). Multiple regimes of constrained chromosome motion are regulated in the interphase Drosophila nucleus. Current Biology 11, 1227–1239.

Vialou, V., Feng, J., Robison, A.J., and Nestler, E.J. (2013). Epigenetic mechanisms of depression and antidepressant action. Annu Rev Pharmacol Toxicol 53, 59–87.

Watson, L.A., and Tsai, L.H. (2017). In the loop: how chromatin topology links genome structure to function in mechanisms underlying learning and memory. Current Opinion in Neurobiology 43, 48–55.

Wiesmeijer, K., Krouwels, I.M., Tanke, H.J., and Dirks, R.W. (2008). Chromatin movement visualized with photoactivable GFP-labeled histone H4. Differentiation 76, 83–90.

Wittmann, M., Queisser, G., Eder, A., Wiegert, J.S., Bengtson, C.P., Hellwig, A., Wittum, G., and Bading, H. (2009). Synaptic Activity Induces Dramatic Changes in the Geometry of the Cell Nucleus: Interplay between Nuclear Structure, Histone H3 Phosphorylation, and Nuclear Calcium Signaling. Journal of Neuroscience 29, 14687–14700.

Yamada, T., Yang, Y., Valnegri, P., Juric, I., Abnousi, A., Markwalter, K.H., Guthrie, A.N., Godec, A., Oldenborg, A., Hu, M., et al. (2019). Sensory experience remodels genome architecture in neural circuit to drive motor learning. Nature 569, 708–713.

Yang, Y., Yamada, T., Hill, K.K., Hemberg, M., Reddy, N.C., Cho, H.Y., Guthrie, A.N., Oldenborg, A., Heiney, S.A., Ohmae, S., et al. (2016). Chromatin remodeling inactivates activity genes and regulates neural coding. Science 353, 300–305.

Yap, E.L., and Greenberg, M.E. (2018). Activity-Regulated Transcription: Bridging the Gap between Neural Activity and Behavior. Neuron 100, 330–348.

Yarrow, J.C., Lechler, T., Li, R., and Mitchison, T.J. (2003). Rapid de-localization of actin leading edge components with BDM treatment. Bmc Cell Biology 4, 5.

Ye, J., Zhao, J., Hoffmann-Rohrer, U., and Grummt, I. (2008). Nuclear myosin I acts in concert with polymeric actin to drive RNA polymerase I transcription. Genes Dev 22, 322–330.

